# Integrated multi-omics reveals common properties underlying stress granule and P-body formation

**DOI:** 10.1101/2020.05.18.102517

**Authors:** Christopher J. Kershaw, Michael G. Nelson, Jennifer Lui, Christian P. Bates, Martin D. Jennings, Simon J. Hubbard, Mark P. Ashe, Chris M. Grant

## Abstract

Non-membrane-bound compartments such as P-bodies (PBs) and stress granules (SGs) play important roles in the regulation of gene expression following environmental stresses. We have systematically determined the protein and mRNA composition of PBs and SGs formed in response to a common stress condition imposed by glucose depletion. We find that high molecular weight (HMW) complexes exist prior to glucose depletion that may act as seeds for the further condensation of proteins forming mature PBs and SGs. Both before and after glucose depletion, these HMW complexes are enriched for proteins containing low complexity and RNA binding domains. The mRNA content of these HMW complexes is enriched for long, structured mRNAs that become more poorly translated following glucose depletion. Many proteins and mRNAs are shared between PBs and SGs including several multivalent RNA binding proteins that may promote condensate interactions during liquid-liquid phase separation. Even where the precise identity of mRNAs and proteins localizing to PBs and SGs is distinct, the mRNAs and proteins share common biophysical and chemical features that likely trigger their phase separation.

## INTRODUCTION

Membrane-bound organelles such as the mitochondria and endoplasmic reticulum provide permanent, tailor-made sub-cellular environments to perform specialized functions within cells, supporting the sequestration of biochemical reactions in a confined and concentrated manner. There is also now increasing evidence that the formation of biological condensates or so-called ‘membrane-less organelles’ act via intracellular phase separation to similarly provide specialized microenvironments, albeit in a more dynamic and environmentally sensitive fashion (Kroschwald, Maharana et al., 2015). These biological condensates are exemplified by a number of ribonucleoprotein (RNP) granules that represent key determinants of mRNA fate in eukaryotic cells with wide-ranging roles in post-transcriptional control (Anderson & Kedersha, 2009).

Two well-studied examples of these RNP granules that exhibit biophysical properties associated with biological condensates are mRNA processing bodies (P-bodies, PBs) and stress granules (SGs) (Anderson & Kedersha, 2009). The localization of mRNAs to PBs and SGs is normally associated with translation repression and they have been accordingly ascribed functions in the degradation and storage of mRNA. SGs and PBs both provide stress-induced microenvironments which share RNA and protein components, and can physically associate with one another. It has been proposed that mRNAs selected for degradation can be passed from SGs to PBs (Kedersha, Stoecklin et al., 2005), although the exact relationship between these different RNP granules remains unclear (Hoyle, Castelli et al., 2007, Kedersha et al., 2005, Shah, Zhang et al., 2013). There is also conflicting data regarding the relationship between the assembly and disassembly of PBs/ SGs: for example, it has been proposed that SG assembly is dependent on PB formation (Buchan, Muhlrad et al., 2008), while it has also been suggested that these condensates arise independently (Hoyle et al., 2007, Shah et al., 2013).

Early evidence suggested that PBs represent sites of mRNA ‘processing’, where an mRNA is decapped and degraded by the 5’–3’ mRNA decay pathway, whereas SGs correspond to sites of mRNA sorting and storage (Lui, Campbell et al., 2010, Sheth & Parker, 2003, Yamasaki & Anderson, 2008). However, more recent studies where PBs have been isolated from mammalian cells indicate that PBs are also involved in mRNA storage (Hubstenberger et al., 2017). In support of an mRNA storage role, mRNAs that are localised to either PBs or SGs can re-enter the pool of mRNAs available for translation after exiting the condensate (Brengues, Teixeira et al., 2005). Such observations further highlight the dynamic and fluid nature of the PBs and SGs within cells.

Generally, RNP granules can either form via the assembly of RNA with protein aggregates, or alternatively, through liquid-liquid phase separation where weak multivalent interactions between multi-domain proteins and RNA form liquid-like droplets in cells (Weber & Brangwynne, 2012). In yeast, PBs appear to form as liquid droplets, whereas SGs resemble more solid protein aggregates (Kroschwald et al., 2015). Further analysis suggests that yeast SGs contain both a more stable core structure encompassed by a phase-separated dynamic outer shell (Jain, Wheeler et al., 2016). Hence, both PBs and SGs are dynamic in nature and appear to rely on ATP-dependent RNA remodelling complexes for their formation and regulation (Hondele, Sachdev et al., 2019, Jain et al., 2016).

Low complexity protein domains and RNA serve as key determinants of the intracellular phase separation necessary for the formation of PBs and SGs (Weber & Brangwynne, 2012). Hence, both PBs and SGs are known to be enriched for proteins containing RNA-binding domains and intrinsically disordered regions (IDRs) (Youn, Dyakov et al., 2019). IDRs are protein domains usually containing stretches of low sequence complexity which have been implicated in the formation of stress-induced RNP granules, as well as in the aggregation of unproductive and cytotoxic protein forms, such as amyloid protein (Gilks, Kedersha et al., 2004). IDRs are generally viewed as having little structure and a high probability of forming amyloidogenic aggregates; they are often referred to as prion-like domains. They have also been ascribed functional roles acting as molecular switches, forming more productive structures when binding with cognate partners such as other proteins or nucleotides (Sandhu & Dash, 2007). Indeed, one attractive possibility is that interactions between multiple IDRs, RBDs and RNA drive PB and SG formation, although our recent studies (Lui, Castelli et al., 2014) suggest pre-existing translation factories can also be remodelled after stress and coalesce with the mRNA decay machinery to seed the formation of bodies.

Given that low complexity protein domains and RNA are major requirements of intracellular phase separation (Weber & Brangwynne, 2012), a fundamental question arises as to how the specificity of composition and function is dictated across different classes of RNA granule in the same cell. The aim of this study was to isolate PBs and SGs following induction by a common stress condition, and then to define any specificity in the attendant transcriptome and proteome of the respective condensates. Here, we integrated a novel quantitative proteomic strategy with immunoprecipitations to isolate and compositionally dissect PBs and SGs after glucose depletion. We find that constituents of these cytoplasmic foci appear to be present as ‘seeds’ prior to stress. Following translation inhibition, these seed elements likely act as a focal point for further protein and RNA condensation. We further show that PBs and SGs are not as discreet as originally hypothesised with several PB protein markers present in SGs and vice versa. Despite differences in the identities of the mRNAs localized to SGs and PBs, we show that the individual mRNA cohorts share common biophysical properties consistent with such properties dictating which mRNAs localize to condensates.

## RESULTS AND DISCUSSION

### Isolation and proteome identification of P-bodies and Stress granules

PBs and SGs are distinct membrane-less protein and RNA condensates that in yeast are both induced by stresses such as glucose starvation. To allow a quantitative assessment of their contents, we used an immuno-affinity approach to isolate PBs and SGs formed in response to glucose depletion. Dcp1p and Pbp1p/Ataxin-2 were chosen as hallmark proteins associated with the two granule types, based on the extensive previous literature that has used these proteins to study and characterize PBs and SGs (Buchan et al., 2008, Hoyle et al., 2007, Sheth & Parker, 2003, Swisher & Parker, 2010). We used strains carrying the myc-tagged markers: Dcp1-myc for PBs, and Pbp1-myc for SGs, to facilitate granule purification. Based on our previous studies, PB and SG formation was induced by 10 or 60 minutes of glucose depletion, respectively (Hoyle et al., 2007, Simpson, Lui et al., 2014).

We first confirmed that the glucose depletion used in these tagged strains causes the well-characterized inhibition of translation initiation that is required for both PB and SG formation (Supplementary Fig. 1). More specifically, across polysome profiles an accumulation of inactive 80S monosomes and depletion of ribosomes from the polysomal portion of the sucrose gradients was observed after glucose depletion (Ashe, De Long et al., 2000). Immunoblot analysis showed that markers for the small and large ribosomal subunits, Rps3 and Rpl35, respectively shift from the polysomal into the monosomal region of the gradients in parallel with the translation inhibition. In contrast, Dcp1 and Pbp1, which are identified across the gradient for unstressed samples, maintain their distribution following glucose depletion. This suggests that after glucose depletion both Dcp1 and Pbp1 are associated with large non-ribosomal protein structures that sediment into sucrose gradients.

PBs and SGs were analysed using label-free mass spectroscopy following the fractionation approach outlined in Fig. 1A. More specifically, whole cell extracts were prepared from Dcp1p-myc and Pbp1p-myc tagged strains in triplicate, with and without glucose starvation; time points were matched to equivalently treated untagged strains. Following formaldehyde cross-linking, cell debris and any unbroken cells were removed via a gentle *1000 x g* centrifugation step to yield a Total fraction (T). An initial centrifugation step (20,000 *x g*) then separated a high molecular weight (HMW) complex (P: pellet fraction) away from supernatant (S). PBs and SGs were isolated from the P-fraction by immunoprecipitation of Dcp1p or Pbp1p, generating ‘unbound’ (U) and ‘elution’ (IP) fractions (Fig. 1A). This fractionation / immunoprecipitation approach was verified by immunoblotting the various fractions for Dcp1-myc and Pbp1-myc (Fig. 1B and C).

**Fig. 1.**
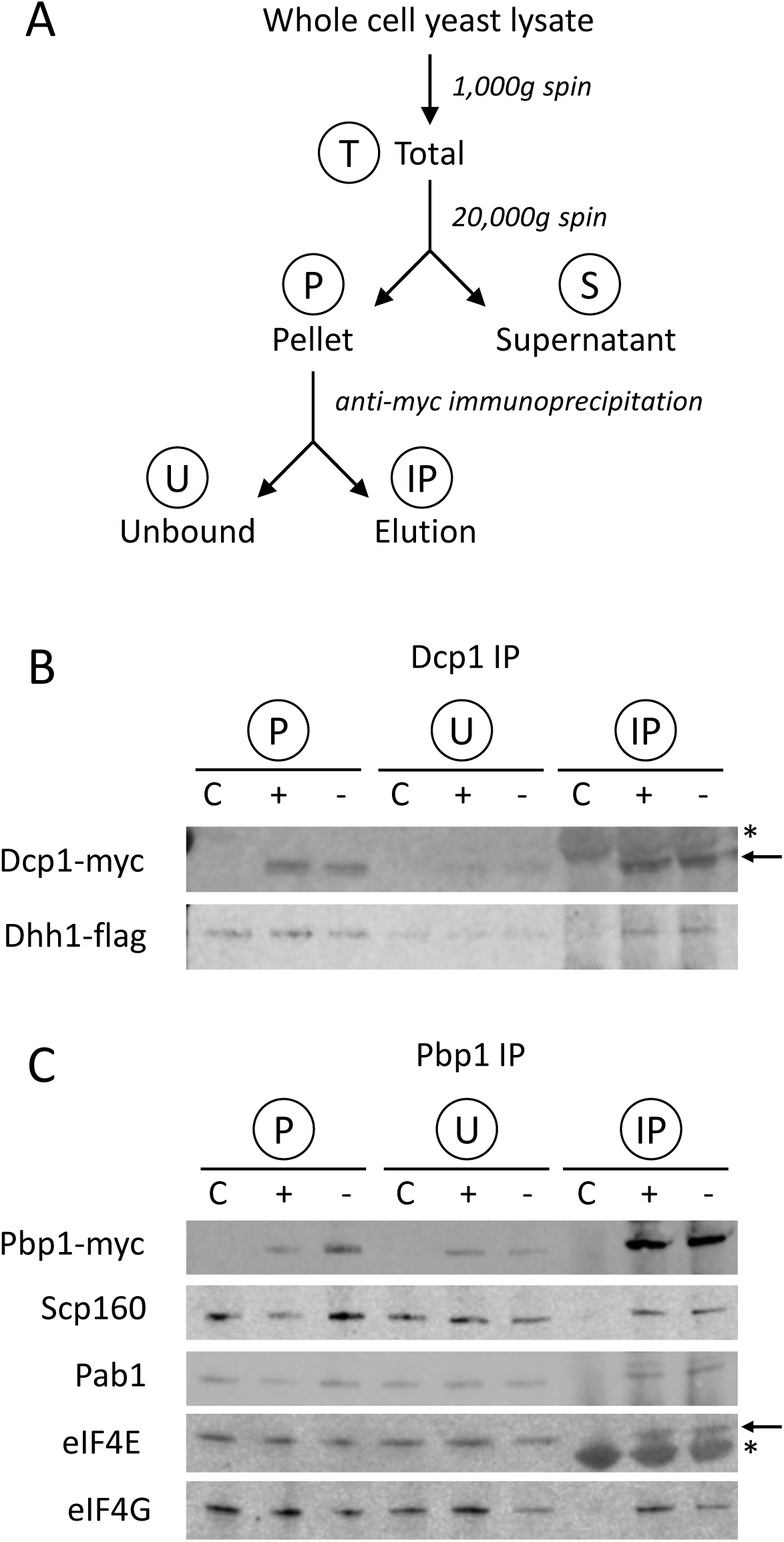
Purification of PBs and SGs. **A.** Schematic of purification process used to isolate PBs and SGs. **B.** Dcp1-myc co-immunoprecipitation of Dhh1-Flag as confirmation of the purification protocol for PBs. Contaminating IgG light chain bands are indicated (asterisk). **C.** Pbp1-myc co-immunoprecipitation of Scp160, Pab1, eIF4E and eIF4G as confirmation of the purification of SGs. Contaminating IgG heavy chain bands are indicated (arrows).

To enable a more nuanced approach to determining the components of PBs and SGs, we adapted the Localisation of Organelle Proteins by Isotope Tagging (LOPIT) technique of Dunkley and colleagues (Dunkley, Watson et al., 2004) to distinguish true condensate components from false positives. A quantitative protein profile is determined by mass spectrometry, composed of signal from across the various separated fractionations, and then used to assign the subcellular localization of proteins by association with known markers. We created a proxy for the separation gradients used in LOPIT using Total, Supernatant, Pellet, Unbound and Elution fractions, then analysed these using label-free mass spectroscopy (Fig.1A). Across all samples, fractions and replicates, 2186 corresponding protein groups were detected at a 1% FDR representing a significant fraction of the yeast proteome. In order to assign proteins to a given condensate, we took an ‘IP-centric’ approach and further considered only proteins detected in at least one tagged IP elution; these fractions should contain all condensate proteins directly or indirectly associated with the marker. Following imputation of missing peptide data, 467 proteins remained, and replicate-averaged quantitative data were split into untreated and glucose deplete sets for further analysis.

To assign proteins to condensates, we clustered the quantitative proteomic data into distinct profiles representing different relative enrichments across the eluted fractions indicated in Fig. 1 (IP, P, S, T and U), using MaxQuant Label-free Quantification (LFQ) intensity (Cox, Hein et al., 2014) as a proxy for protein abundance. Since proteins are not necessarily exclusive to a single condensate type and can move between them, we used a fuzzy clustering approach to support membership of multiple clusters. We combined the MaxQuant LFQ signals from all fractions and in different conditions into a single vector with either 15 (untreated) or 20 (glucose depleted) values as input for the Mfuzz clustering tool (Kumar & M, 2007). Clusters were determined experimentally to explicitly separate the two condensate marker proteins Dcp1p and Pbp1p into individual clusters, creating ten clusters for the untreated samples (Supplementary Fig. 2A and Supplementary Table 1) and 11 clusters for glucose depleted samples (Supplementary Fig. 2B and Supplementary Table 2).

Presence in one of the two condensate types was inferred from membership of the cluster containing the representative tagged bait protein, Dcp1p or Pbp1p, respectively (Fig. 2A). The Mfuzz algorithm assigns a membership score to each protein for each cluster and hence initially, proteins are assigned to the cluster to which they have the highest membership score. The Dcp1p and Pbp1p immunoprecipitates derived from unstressed cells are referred to as ‘pre-P-bodies’ (pre-PBs) (Fig. 2A, Cluster 1) and ‘pre-stress granules’ (pre-SGs) (Fig. 2A, Cluster 4), respectively. After glucose depletion, immunoprecipitates are referred to as PBs (Fig. 2A, Cluster 7) and SGs (Fig. 2A, Cluster 11).

**Fig. 2.**
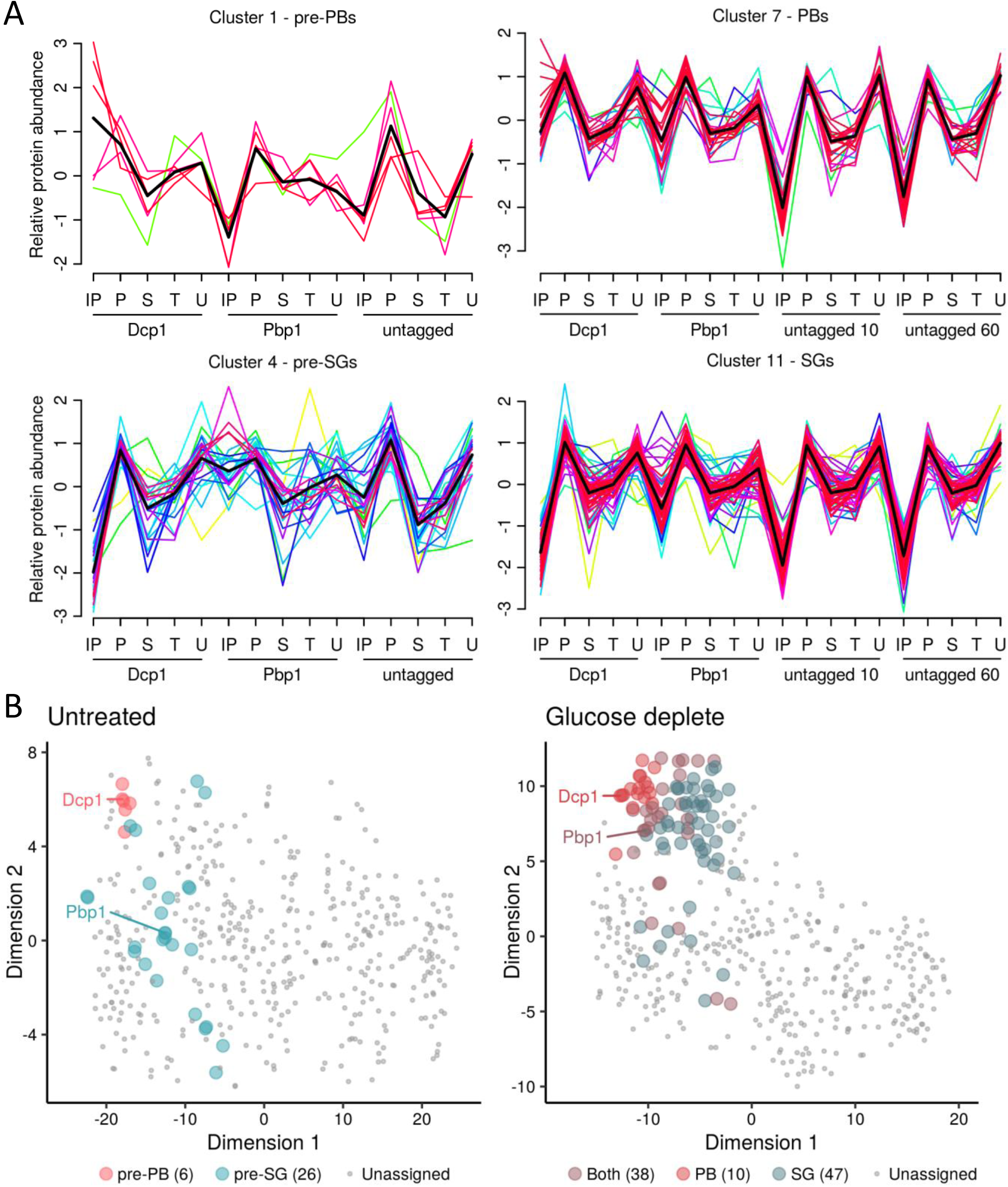
Identification of protein components of PBs and SGs using clustering. **A.** Known PB and SG components were used to designate clusters representing pre-PBs (Cluster 1), PBs (Cluster 7), pre-SGs (Cluster 4), and SGs (Cluster 11). Lines are coloured by how well each protein correlates with the cluster and a black line represents the average of all data within the cluster. **B.** PCA analysis of proteomic data under untreated and glucose depleted conditions. The t-SNE dimensionality reduction technique is used to show the relationship between the proteins as described by their quantitative proteomic profiles across all fractions, coloured by their cluster membership.

As can be seen in Fig. 2A, for example, six proteins in total are members of the pre-PB cluster, displaying coherent, common elution profiles across all 15 fractions submitted for MS analysis. In simple terms, these proteins are over-represented in the IP fractions for Dcp1p-myc, but not in the corresponding IPs for Pbp1p or the untagged control. A more nuanced interpretation is that these profiles are more complex than a simple ‘standard’ differential expression approach comparing the tagged IP *versus* untagged IP or similar pairwise analyses. By considering a common relative abundance throughout *all* 15 fractions we are better able to distinguish truly interacting proteins that display similar properties to the marker protein, and also exclude some high abundance ‘false positives’ that are commonly represented in immunoprecipitation experiments (Mellacheruvu, Wright et al., 2013). For example, the clustering approach allowed us to identify proteins that have been previously been shown to be in PBs (Lsm1, Scd6 and Pop2) and SGs (eIF4G1, eIF4G2 and Ygr250c) that are not captured from the differential expression approach comparing the tagged IP *versus* an untagged control (Supplementary Tables 3 and 4).

The fuzzy clustering approach has an additional advantage; since each protein has a membership score to each cluster, we determined the proteins whose two highest membership scores were to the clusters containing the two condensate marker proteins, and defined these as overlapping proteins with membership to both PBs and SGs. There were no such proteins in unstressed conditions, but 38 in stressed conditions satisfied this criteria, suggesting overlapping components for these two cluster types. This data is represented in Fig. 2B where the t-Distributed Stochastic Neighbour Embedding (t-SNE) dimensionality reduction technique shows the relationship between the proteins from their quantitative proteomic profiles, coloured by their cluster membership. In the untreated plot there is good separation between the two pre-granule bodies, whilst in glucose deplete conditions the protein points overlap more – indeed many of the PB assigned proteins are also members of the SG cluster and vice versa.

### P-body and Stress granule ‘seeds’ exist prior to glucose starvation

In the untreated condition, we identified six pre-PB proteins, including Dcp1, Dcp2 and Edc3; and 26 pre-SG proteins (Fig. 3A and B, Table 1 and 2). In the glucose deplete conditions, these lists expand to 47 proteins in PBs and 85 proteins in SGs suggesting that glucose depletion causes condensation of proteins to the Dcp1p and Pbp1 ‘seeds’ that enlarge to form PBs and SGs (Fig. 3A and B). Interestingly, not all of the proteins assigned to pre-PBs and pre-SGs were present following glucose starvation suggesting that, not only does condensation of new proteins occur during stress, but some pre-stress proteins are lost and there is remodelling of the pre-PBs and pre-SGs.

**Fig. 3.**
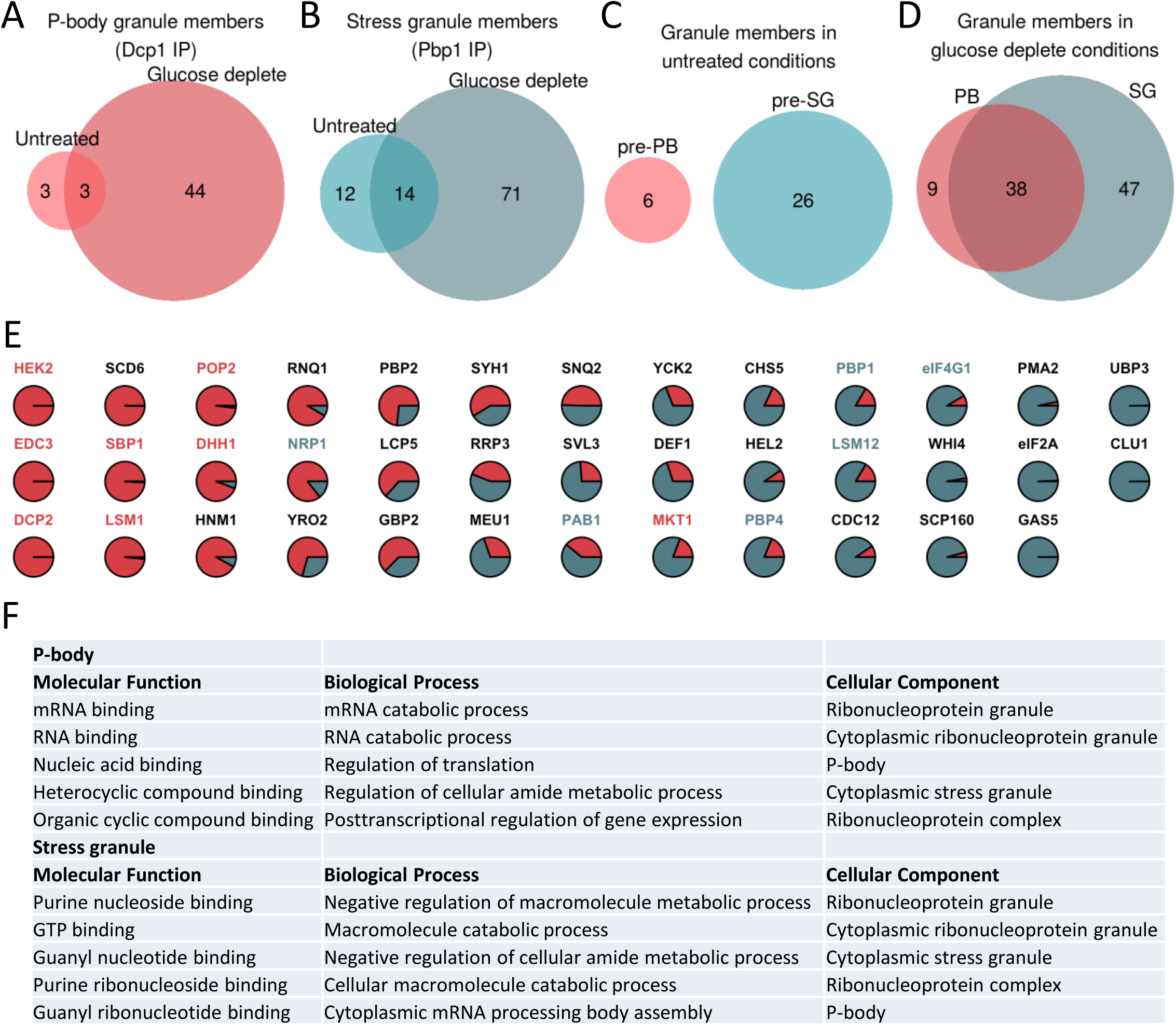
PB and SG seeds exist prior to stress and act as sites of protein condensation driven by protein disorder. **A.** Euler diagram comparing protein components of pre-PBs and PBs. **B.** Euler diagram comparing protein components of pre-SGs and SGs. **C.** Euler diagram comparing protein components of pre-PBs and pre-SGs. **D.** Euler diagram comparing protein components of PBs and SGs **E.** Cluster membership scores for proteins assigned to PBs and SGs. PB (cluster 7) membership coloured red and SG (cluster 11) membership coloured green. **F.** Functional categorisation of proteins present in PBs and SGs following glucose depletion. The top five categories (Molecular function, Biological process and Cellular component) are shown for PBs and SGs taken from the complete analysis shown in Supplementary Fig. 3.

**Table 1.**
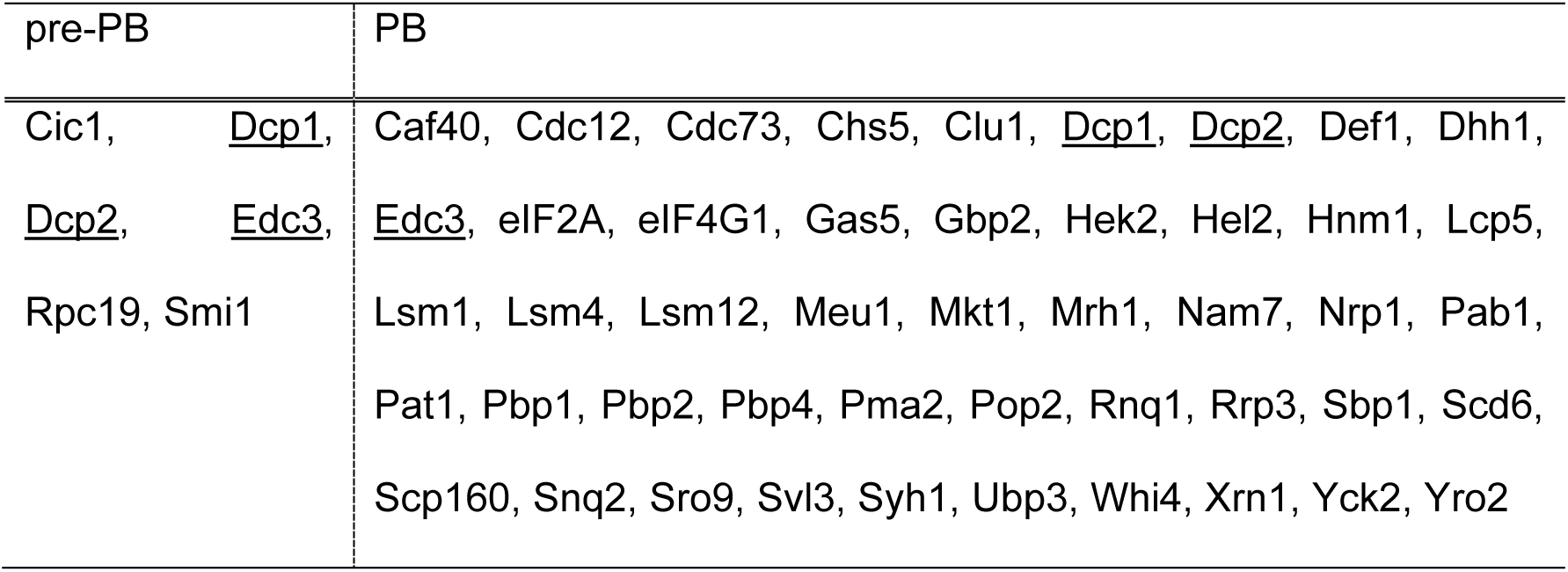
Proteins interacting with Dcp1p-myc with and without glucose.

| pre-PB | PB |
| --- | --- |
| Cic1, <u>Dcp1</u>,<br><u>Dcp2</u>, <u>Edc3</u>,<br>Rpc19, Smi1 | Caf40, Cdc12, Cdc73, Chs5, Clu1, <u>Dcp1</u>, <u>Dcp2</u>, Def1, Dhh1,<br><u>Edc3</u>, eIF2A, eIF4G1, Gas5, Gbp2, Hek2, Hel2, Hnm1, Lcp5,<br>Lsm1, Lsm4, Lsm12, Meu1, Mkt1, Mrh1, Nam7, Nrp1, Pab1,<br>Pat1, Pbp1, Pbp2, Pbp4, Pma2, Pop2, Rnq1, Rrp3, Sbp1, Scd6,<br>Scp160, Snq2, Sro9, Svl3, Syh1, Ubp3, Whi4, Xrn1, Yck2, Yro2 |

**Table 2.** Proteins interacting with Pbp1p-myc with and without glucose.

| pre-SG | SG |
| --- | --- |
| Brx1, <u>Bsp1</u>, <u>Bud7</u>,<br>Cdc73, Enp2, Gar1,<br><u>Hnm1</u>, <u>Lsm12</u>,<br><u>Mkt1</u>, Nma1, Nop1,<br>Osh6, <u>Pbp1</u>, <u>Pbp2</u>,<br><u>Pbp4</u>, <u>Puf3</u>, Puf6,<br>Rpf2, Rpb2, Rpn5,<br><u>Rrb1</u>, <u>Ssh1</u>, <u>Svl3</u>,<br><u>Syh1</u>, <u>Tat1</u>, Utp22, | Bfr1, Bre5, <u>Bsp1</u>, <u>Bud7</u>, Cdc12, Cdc39, Chs5, Cic1, Clu1,<br>Cop1, Coy1, Dcp2, Ded1, Def1, Dhh1, Ecm32, Ecm33, Edc3,<br>eIF2A, eIF3g, eIF3i, eIF4G1, eIF4G2, Fun12, Gas1, Gas5,<br>Gbp2, Hek2, Hel2, <u>Hnm1</u>, Hxt1, Ist2, Lcp5, Lsm1, <u>Lsm12</u>,<br>Map1, Meu1, Mis1, <u>Mkt1</u>, Mrn1, Nat1, Nce102, Nop12, Nop4,<br>Nrp1, Pab1, <u>Pbp1</u>, <u>Pbp2</u>, <u>Pbp4</u>, Phb1, Pil1, Pma1, Pma2,<br>Pop2, Pub1, <u>Puf3</u>, Puf4, Rbg1, Rho1, Rie1, Rnq1, <u>Rrb1</u>,<br>Rrp3, Sbp1, Scd6, Scp160, Sec1, Sec27, Sgn1, Snq2, Srp54,<br>Srp68, <u>Ssh1</u>, Sui3, Sup35, Sur7, <u>Svl3</u>, <u>Syh1</u>, <u>Tat1</u>, Ubp3,<br>Whi2, Whi4, Yck2, Yro2 |

Underlined proteins are those present in both pre-PBs and PBs or pre-SGs and SGs.

A number of studies have highlighted that PBs and SGs have different components, leading to the suggestion that they are functionally distinct (Youn et al., 2019). SGs act to store mRNAs that can re-enter the translationally active pool of mRNAs, whilst PBs were originally thought to play a key role in mRNA decay. Consistent with this view, the protein components of pre-PBs and pre-SGs show no overlap under unstressed conditions (Fig. 3C). After glucose depletion however, 38 proteins are present in both PBs and SGs suggesting that PBs and SGs are not as compositionally distinct as expected (Fig. 3D). These data are consistent with a complex pattern where proteins are distributed across different pools in PBs and SGs, rather than being unique to particular condensates. In fact, the 38 ‘common’ proteins include several proteins that have previously been used by numerous labs (including ourselves) to specifically localise PBs (Edc3p, Dcp2, Dhh1p, Sbp1p, Lsm1p, Hek2p, Mkt1p, Pop2p) or SGs (Pab1p, Pbp1, eIF4G1, Pbp4p, Lsm12p, Nrp1p) (Buchan et al., 2008, Jain et al., 2016, Lewis, Broman et al., 2014, Mitchell, Jain et al., 2013, Reijns, Alexander et al., 2008, Swisher & Parker, 2010), respectively. By examining cluster membership scores for the proteins assigned to both condensates, it is possible to observe whether they associate more strongly with either PBs or SGs. Of these previously used condensate markers, all cluster more closely with the granule type they were used as a marker for other than Nrp1p and Mkt1p (Fig. 3E). This situation is entirely analogous to what has been described for SG and PB proteomics in mammalian cells (Youn et al., 2019). Therefore, the distinction between these condensates in both yeast and mammals is not absolute, reflecting possible interactions between PBs and SGs. It also seems likely that the localization of individual proteins depends upon numerous factors including the nature of the stress and the proteomic context of cells pre-stress.

PBs might be expected to contain proteins involved in translational repression, mRNA decapping and 5’ to 3’ exonuclease activity given their proposed role in mRNA decay, whereas, SGs might be expected to contain translation factors and translation regulatory factors in accordance with their proposed role in mRNA and translation factor storage. Given the small numbers of proteins identified in pre-PBs and Pre-SGs, only those proteins present in PBs and SGs formed after stress were examined for enrichment of Gene Ontology (GO) terms (Supplementary Fig. 3). The top five most significant categories identified in each case are also highlighted in Fig. 3F. This analysis confirmed that both PBs and SGs are enriched for the RNP granule components category, as well as enrichments in proteins from both the P-body and stress granule categories, consistent with our observed overlap between PBs and SGs. Furthermore, both condensates are enriched in broad GO categories including mRNA catabolism, deadenylation-dependent decay, deadenylation-dependent decapping of mRNA, regulation of translation and translation initiation, further suggesting that these condensates interact and contain overlapping components.

The proteome of SGs induced by sodium azide stress has been studied previously (Jain et al., 2016). We observed a modest but significant overlap of 23 proteins identified in comparison to sodium azide induced SGs (Supplementary Fig. 4A). The core sodium azide SG proteome was extended to include additional SG proteins that could not be confirmed by microscopy or had only been observed *in vitro* (Jain et al., 2016). However, the overlap between the glucose-depletion induced SG proteome and this extended sodium azide SG proteome only increases to 25 proteins (Supplementary Fig. 4B) suggesting that SGs induced under glucose depletion and sodium azide stress have different protein complements. The study by Jain *et al*. also generated a PB proteome by literature mining. Comparing this PB proteome to our glucose depletion PB data set identified a small number (16) of common proteins with much larger numbers of proteins unique to each PB data set (Supplementary Fig. 4C). Not all of the experiments that were used to create the Jain *et al*. PB proteome used glucose depletion to drive PB formation suggesting that certain PB components are localised to these foci in a stress specific manner. Intriguingly these data show that the similarities between SGs and PBs formed after glucose depletion are more striking than the similarities between SGs formed under different conditions, or PBs formed under different conditions. These results highlight the remarkably intricate stress specific composition of both SGs and PBs.

### P-bodies and stress granules are enriched for proteins containing regions of low complexity and under-enriched for proteins with regions of hydrophobicity

Given the differences in protein composition of condensates identified under different stress conditions, we next asked whether their protein components share similar biophysical properties. For instance, regions of low complexity within proteins can increase their capacity to phase separate (Zhou, Nguemaha et al., 2018). We assessed the protein subsets present in our sample preparations for both intrinsically disordered regions and the proportion of disordered residues. Strikingly, the probability of containing an intrinsically disordered region is significantly higher in proteins present across our immunoprecipitated complexes when compared to the background proteome (Fig. 4A). Even within these data the probability of containing intrinsically disordered regions is higher in SGs relative to pre-SGs suggesting that the presence of disordered protein regions correlates with the formation of larger condensates.

**Fig. 4.**
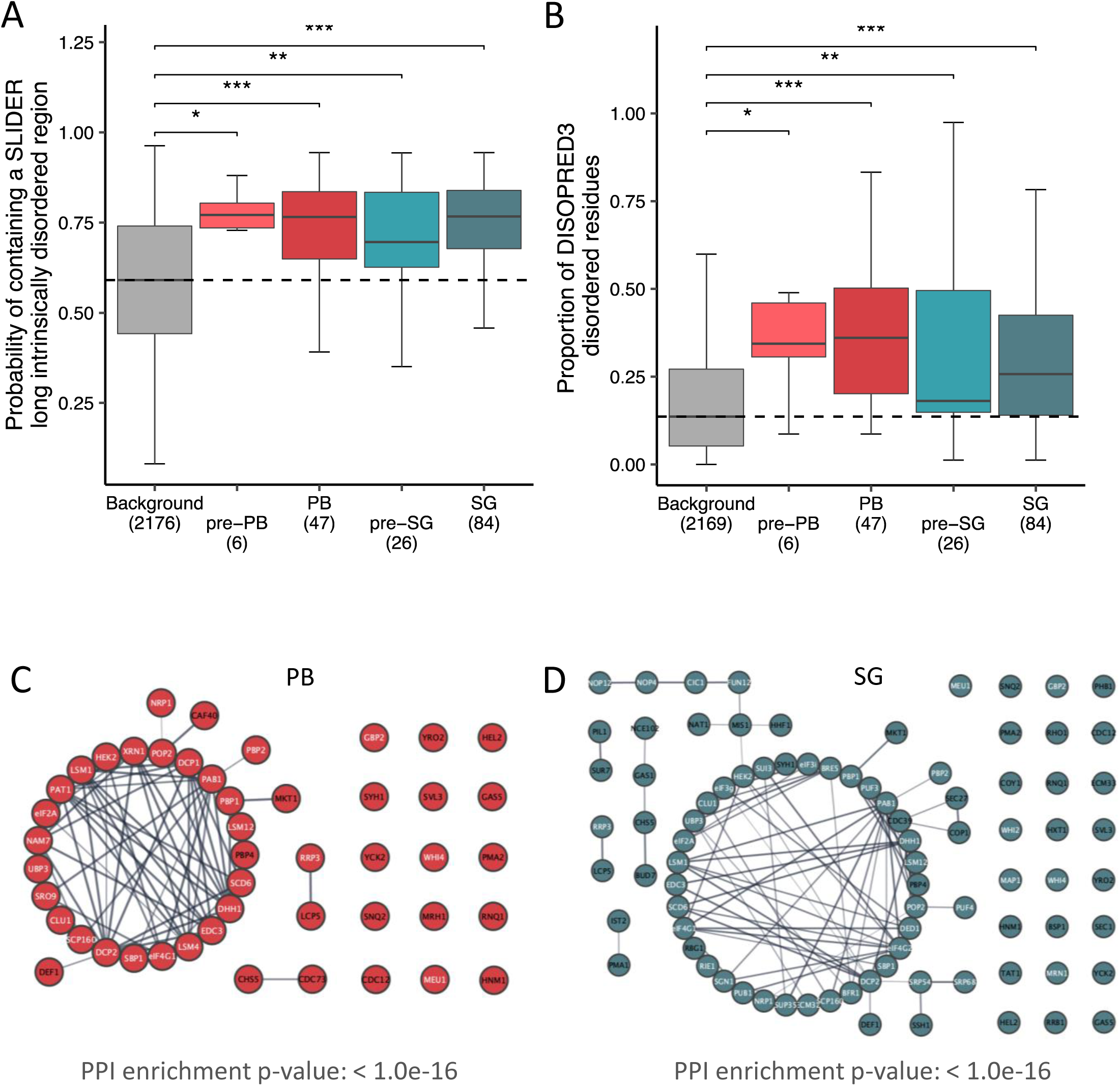
PBs and SGs are enriched for proteins containing regions of low complexity and for proteins with significantly enriched interactomes. **A.** Box-plots comparing the probability of a condensate protein containing a long intrinsically disordered region (determined by SLIDER) with a background protein set (all proteins seen in the mass spectroscopy analysis). Black dashed lines represent the median of the background dataset. **B.** Box-plots comparing the probability of a condensate protein containing disordered residues (determined by DISOPRED3). **C.** Analysis of the network of predicted protein interactions (PPI) for PBs focussing on direct physical interactions. Each node represents a protein identified as a member of the PB cluster and previously identified protein:protein interactions are indicated by lines between the nodes. Proteins labelled in white are those that have an RNA binding GO annotation. **D.** Analysis of the PPI for SGs as for panel C.

A similar situation is observed when the proportion of disordered residues per protein is evaluated (Fig. 4B), with significant enrichment across all the immunoprecipitated complexes and higher values in SGs relative to pre-SGs. As a control, the same analysis was performed on the proteins identified in all the fractions collected during condensate preparation. This analysis showed that there is no enrichment for disordered proteins in any of these samples (Supplementary Fig. 5A). In fact, the proportion of disordered residues is significantly reduced in Elution fractions suggesting that the analysis that we have performed to identify PB and SG components, based upon their profile across purification samples relative to known SG/PB components, specifically enriches disordered proteins from a pool of ordered proteins. Furthermore, the average protein length remains the same across all fractions suggesting that the centrifugation steps used to enrich condensates and subsequent immunoprecipitation is specific and not simply enriching for longer, and thus heavier proteins (Supplementary Fig. 5B).

We analysed the PB and SG protein components further to test whether other biochemical and structural properties characterize the localization of proteins to condensates. This analysis revealed that proteins containing bulky, aromatic and hydrophobic amino acid regions are under-enriched in all of the immunoprecipitates (Supplementary Fig. 5C). Conversely, proteins containing exposed, flexible amino acid regions and regions of high solvation propensity are enriched in PBs and SGs. These data are consistent with the understanding that PBs and SGs are aqueous condensates that arise due to phase separation in the cytoplasm.

### Significantly enriched protein interactomes are identified in P-body and stress granule proteomes

Analysis of proteins components of PBs (Fig. 4C) and SGs (Fig. 4D) following glucose depletion revealed a strong and significant enrichment for known protein-protein interactions, focussing on directly observed physical interactions. In both cases the protein sets form tight interconnected protein groups with many previously characterised molecular interactions, consistent with our purification capturing a true representation of the cognate protein components present in these condensates. The proteins in the resulting networks are also significantly enriched for proteins with RNA binding activity (Proteins labelled in white in Fig. 4C and D) consistent with previous data suggesting that these condensates are enriched for proteins containing RNA-binding domains (RBDs) (Jain et al., 2016, Wang, Schmich et al., 2018, Youn et al., 2019). This observation highlights the important role that RNA plays in the formation and dynamics of intracellular phase separated condensates (Ivanov, Kedersha et al., 2019). Therefore, the next step was to examine the finite RNA composition of PBs and SGs.

### RNA presence in condensates is dependent on key physical characteristics

RNA plays an important role in the formation of PBs and SGs (Hoyle et al., 2007, Hubstenberger, Courel et al., 2017, Lui et al., 2014, Wang et al., 2018) and even has the ability to condense in the absence of protein (Van Treeck, Protter et al., 2018). To identify component mRNAs before and during PB and SG formation, RNA was isolated from condensates using the same fractionation strategy as for our protein analysis. In the absence of established marker mRNAs for PBs and SGs, we used a traditional RIP-Seq approach to enrich for Pbp1p and Dcp1p associated mRNAs and compared to a total RNA prep. Samples were assessed in triplicate from unstressed or stressed cells using RNA-Seq, then expressed as an enrichment in the immunoprecipitates relative to total RNA.

Dcp1p RIP-Seq on unstressed cells identified 1717 significantly enriched RNAs which decreased to 1433 RNAs after glucose depletion. Pbp1p enriched 2116 RNAs under unstressed conditions compared with 1591 RNAs after glucose starvation. In keeping with the proteomic interaction between SGs and PBs there was a large overlap in the mRNAs shared between pre-PBs and pre-SGs, and between PBs and SGs (Supplementary Fig. 6A and B). Equally, pairwise comparisons revealed significant similarity between the stressed and unstressed data-sets especially for the Dcp1p-associated mRNAs (Fig. 5A; R^2^ = 0.62) with weaker correlation for the Pbp1-associated RNAs (Fig. 5B; R^2^ = 0.32). Thus, comparing the stress versus unstressed experiments there is a greater correlation for Dcp1p RIP-Seq datasets than for the Pbp1p RIP-Seq datasets. This suggests there is a more significant remodelling of the RNA content of stress granules after stress as although a core set of 1067 remain associated with Pbp1p, 1049 are lost and 524 become significantly enriched in the IP. This effect is notably reduced in P-bodies (Fig. 5C & D). This change is broadly consistent with the proteomics data which also points to a more significant remodelling of stress granules.

**Fig. 5.**
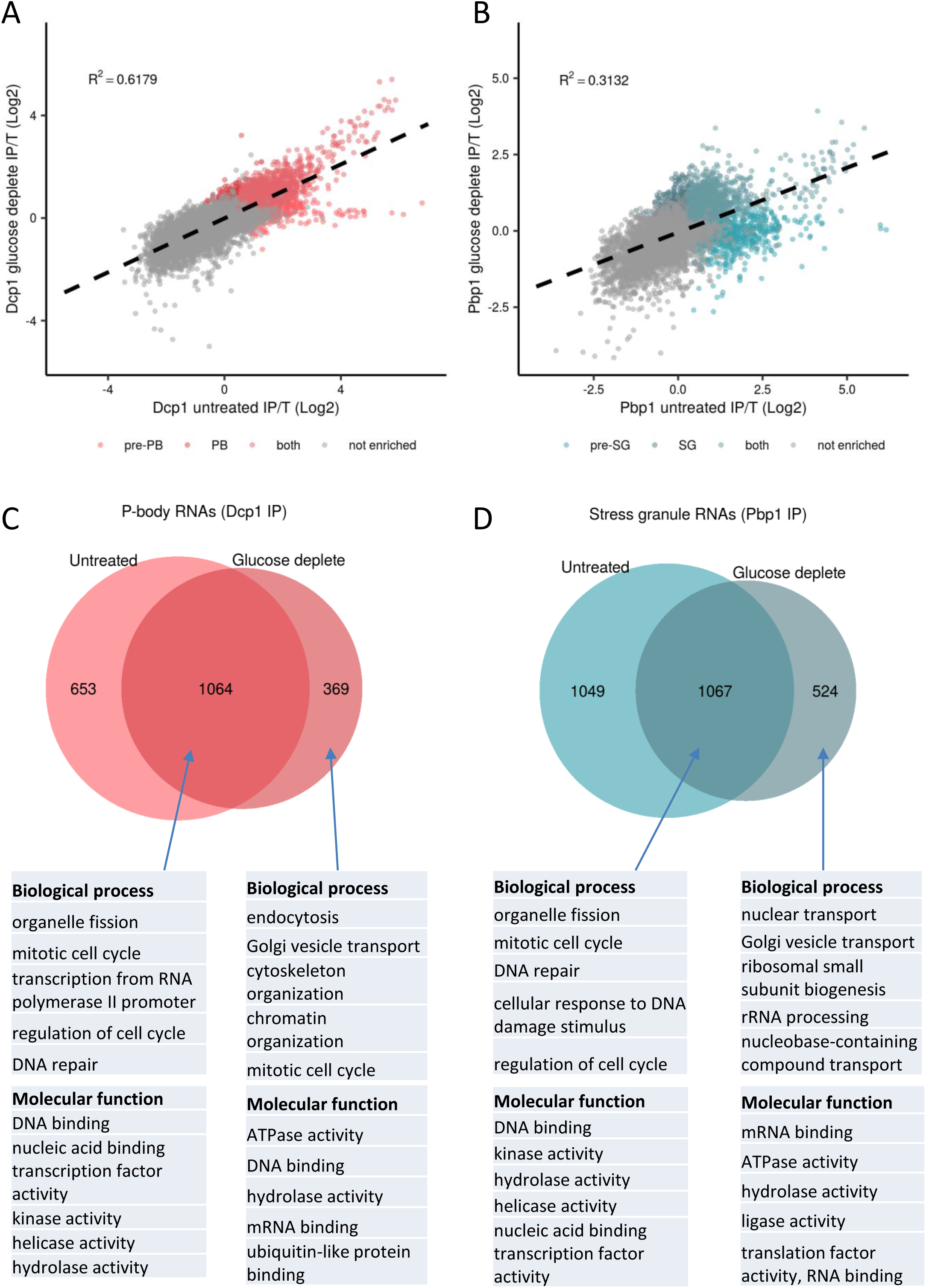
Identification of RNAs isolated from PBs and SGs. **A.** An XY scatterplot comparing Log_2_ fold enrichment of IP/T for Dcp1p-associated mRNAs under untreated conditions or conditions of glucose depletion. **B.** An XY scatterplot comparing Log_2_ fold enrichment of IP/T for Pbp1p-associated mRNAs under untreated conditions or conditions of glucose depletion. **C.** Euler diagram comparing mRNA components of pre-PBs and PBs. The top five GO categories (Molecular function, Biological process) are shown taken from the complete analysis in Supplementary Fig. 7. **D.** Euler diagram comparing mRNA components of pre-SGs and SGs. The top five GO categories (Molecular function, Biological process) are shown taken from the complete analysis in Supplementary Fig. 7.

We also tested for significant GO enrichment of the various RIP-Seq mRNA datasets (Supplementary Fig. 7). For this analysis, the mRNAs were split into various pools: 1. mRNAs that are *uniquely* associated with pre-PBs or pre-SGs (unstressed conditions), 2. mRNAs that *uniquely* associate with PBs or SGs (after glucose depletion) and 3. mRNAs that are identified with both pre-PBs and PBs, or with pre-SGs and SGs. The top five most significant categories from this analysis are highlighted in Fig. 5C and D. Although large numbers of mRNAs were associated uniquely with pre-PBs and pre-SGs, no functional enrichments were observed in the function of the proteins encoded by these mRNAs. This suggests that the mRNAs uniquely localised to pre-PBs and pre-SGs are not particularly co-ordinated in terms of function.

Interestingly, substantial overlap was identified across the enriched mRNA functional categories for the sets of mRNAs that are common to both pre-PBs and PBs, or both pre-SGs and SGs. For instance, mRNAs encoding proteins with DNA and nucleic acid binding activities and proteins affecting processes including organelle fission, mitotic cell cycle and DNA repair are all enriched. These data suggest that transcripts encoding proteins associated with the broad regulation of cellular activities such as the cell cycle are found as granule associated and the specificity between SG or PB localisation does not appear particularly critical. This fits with mammalian cell studies where these types of mRNA were identified as stored in PBs, and suggests that in yeast such storage can occur in either PBs or SGs. In contrast, the mRNAs unique to PBs and SGs encode proteins affecting distinct biological processes. More specifically, PBs are enriched for mRNAs that encode proteins involved in endocytosis, cytoskeleton organization and chromatin organization, whereas, SGs are enriched for mRNAs that encode proteins involved in nuclear transport, rRNA processing and ribosomal subunit biogenesis. These data indicate that while many mRNAs are shared across PBs and SGs, the uniquely localized mRNAs will lead to an impact on different cellular functions and processes.

A previous study assessed the mRNA content of SGs induced by sodium azide stress (Khong, Matheny et al., 2017). Although similar, there is only modest correlation between our four datasets and these data (R^2^ < 0.5) and in fact the glucose-depletion induced SG data-set shows the lowest similarity to the sodium azide SG dataset (Supplementary Fig. 6C - K). This suggests that the identity of mRNAs present in PBs and SGs is particularly dependent on the stress used to induce the condensates, and further highlights the possibility that localization of distinct mRNAs to condensates is part of a stress-specific mRNA rationalization process in cells.

### P-bodies and stress granules are enriched for longer, more highly structured mRNAs

Although the specific mRNAs identified in glucose-depletion induced PBs and SGs are largely different to those seen in sodium azide-induced SGs, they share similar properties. Sodium azide-induced SGs were reported to be enriched for mRNAs with longer ORF lengths (Khong et al., 2017) and mRNAs localizing to our complexes are also significantly longer when compared with mRNA length across the transcriptome (Fig. 6A). This effect is predominantly attributable to an increase in overall CDS length, although the length of both 5’ and 3’ UTRs were also increased for localized mRNAs (Supplementary Fig. 8A-E). The mRNAs that localize to PBs or SGs are also significantly longer than those that localize to pre-PBs or pre-SGs (Fig. 6A) and this difference is again predominantly due to a variation in the length of the CDS (Supplementary Fig. 8A-E).

**Fig. 6.**
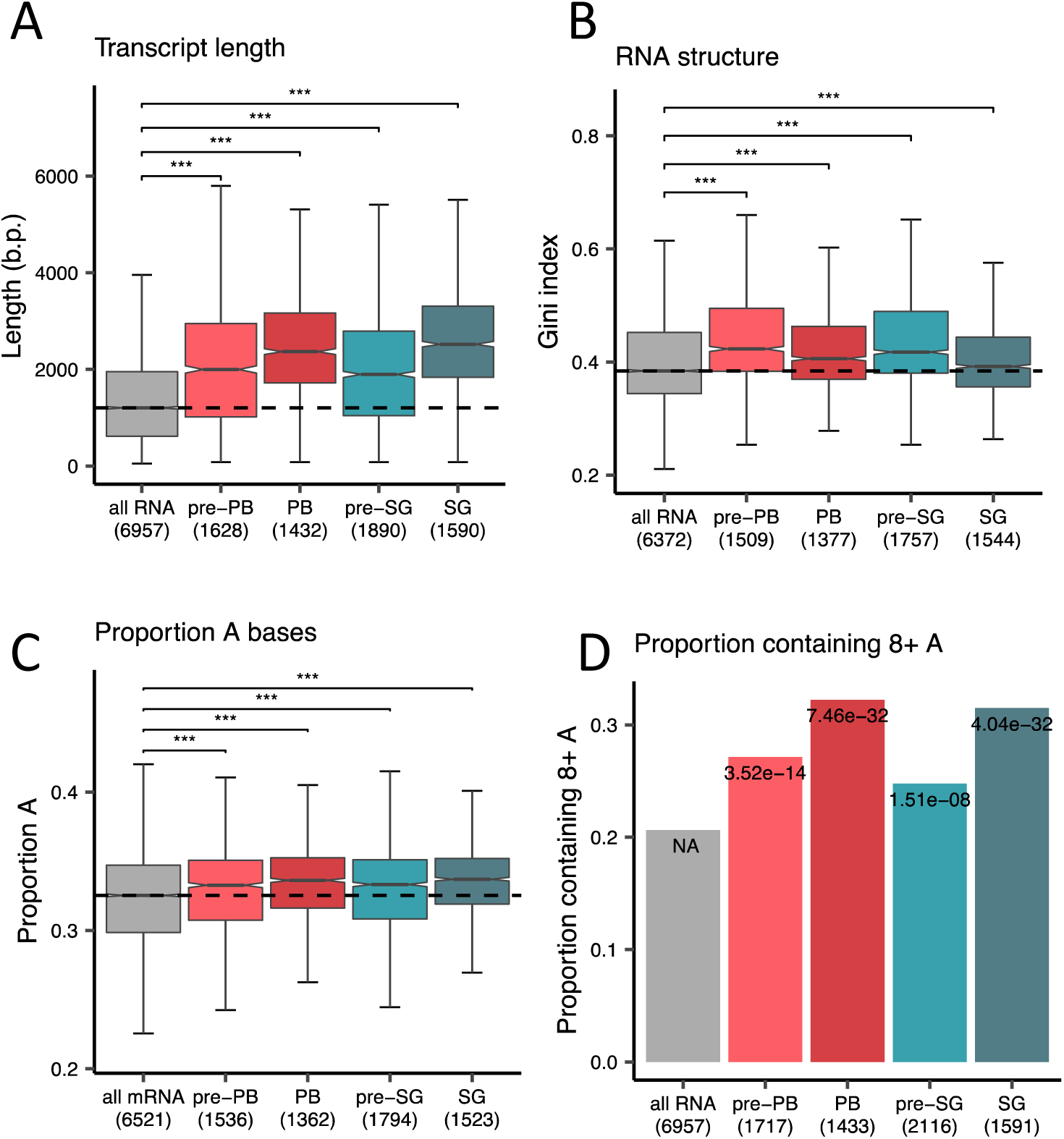
Long, structured mRNAs enriched for adenosine localize to PBs and SGs. **A.** Box plots comparing transcript lengths of mRNAs enriched in pre-PBs, PBs, pre-SGs and SGs with all mRNAs from the transcriptome as a background control. **B.** Box plots comparing RNA secondary structure (Rouskin et al., 2014) of mRNAs enriched in pre-PBs, PBs, pre-SGs and SGs with all mRNAs. **C.** Box plots comparing adenosine content of mRNAs enriched in pre-PBs, PBs, pre-SGs and SGs with all mRNAs. **D.** Box plots comparing polyA tract content (defined as runs of eight or more adenosine residues) of mRNAs enriched in pre-PBs, PBs, pre-SGs and SGs with all mRNAs.

The mRNAs from the RIP-Seq experiments were further examined to determine whether any other biochemical or physical characteristics might account for their condensate localization. RNA has been shown to phase separate *in vitro* (Van Treeck et al. 2018) and long structured RNAs have a propensity to self-assemble into condensates. Using a transcriptome-wide dataset for RNA secondary structure (Rouskin, Zubradt et al., 2014), we found that mRNAs identified in pre-PBs, pre-SGs, PBs and SGs, are on average significantly more structured than the general transcriptome (Fig. 6B). Furthermore, the mRNAs present in PBs and SGs are less structured relative to their pre-PB and pre-SG seeds. Taken together, these data indicate that longer, more structured mRNAs are generally enriched in the condensates, and during glucose depletion, there is a shift to longer, less-structured mRNAs during PB and SG formation from pre-PBs and pre-SGs, respectively.

To explore this further, AU and CG nucleotide content was examined as a proxy for secondary structure in PBs and SGs relative to pre-PBs and pre-SGs. The coding regions of mRNAs in the condensates are enriched for AU compared to the transcriptome average (Supplementary Fig. 8F), which parallels observations made for sodium azide-induced SGs (Khong et al., 2017). Moreover, the AU enrichment for SGs, PBs, pre-PBs and pre-SGs arises predominantly from an increase in adenosine content (Fig. 6C and Supplementary Fig. 8G). Interestingly, this increased adenosine content may be explained by an enrichment for polyA tracts (Fig. 6D). Once again and consistent with the observations above for secondary structure, mRNAs detected in PBs and SGs are more enriched for adenosine residues and polyA tracts than those in pre-PBs and pre-SGs.

Taken together, these data indicate that although different mRNAs localize to PBs and SGs, in what appears to be a stress dependent-manner, they share common biophysical properties which distinguish their condensate localization from the general mRNA pool.

### The translational efficiency of mRNAs localised to P-bodies and stress granules is reduced after glucose starvation

Given that both SGs and PBs are sites where translationally repressed mRNAs become localised (Kedersha, Chen et al., 2002, Teixeira, Sheth et al., 2005), we performed a ribosome profiling analysis (McGlincy & Ingolia, 2017) to directly assess the translation efficiency (TE) of the mRNAs identified in pre-PBs, pre-SGs, PBs and SGs. As a first step, we evaluated the TE of these mRNAs under active growth (non-stress) conditions. Those mRNAs that uniquely associate with PBs or SGs after glucose depletion had the highest translation efficiency when measured in unstressed cells, whereas the mRNAs uniquely associated with pre-PBs or pre-SGs had the lowest TE under these conditions (Supplementary Fig. 9A and B). This indicates that highly translated mRNAs are less enriched in pre-PBs and pre-SGs, yet after the large-scale translation inhibition caused by glucose depletion, many previously highly translated mRNAs are now found in PBs and SGs.

Further confirmation of the inverse relationship between SG/PB localisation and translation level comes from the analysis of translation efficiencies in cells after glucose depletion. The TE data used for our initial analysis was determined in cells grown under conditions where glucose was available in the growth media. To extend this analysis we used matched TE data obtained from cells following 10 min glucose depletion conditions. The mRNAs uniquely associated with Dcp1p and Pbp1p under unstressed conditions showed higher translation efficiencies following glucose depletion, when they were no longer apparently significantly associated with Dcp1p or Pbp1p (Fig. 7). Conversely, the TEs of those mRNAs uniquely associated with Dcp1p or Pbp1p following glucose starvation reversed this trend, and were found to be reduced following glucose depletion, suggesting that the localisation of these mRNAs to PBs and SGs correlates with a reduction in translation (Fig. 7). These trends are also consistent with theoretical predictions of protein translation based around codon usage and tRNA availability (Supplementary Fig. 9C).

**Fig. 7.**
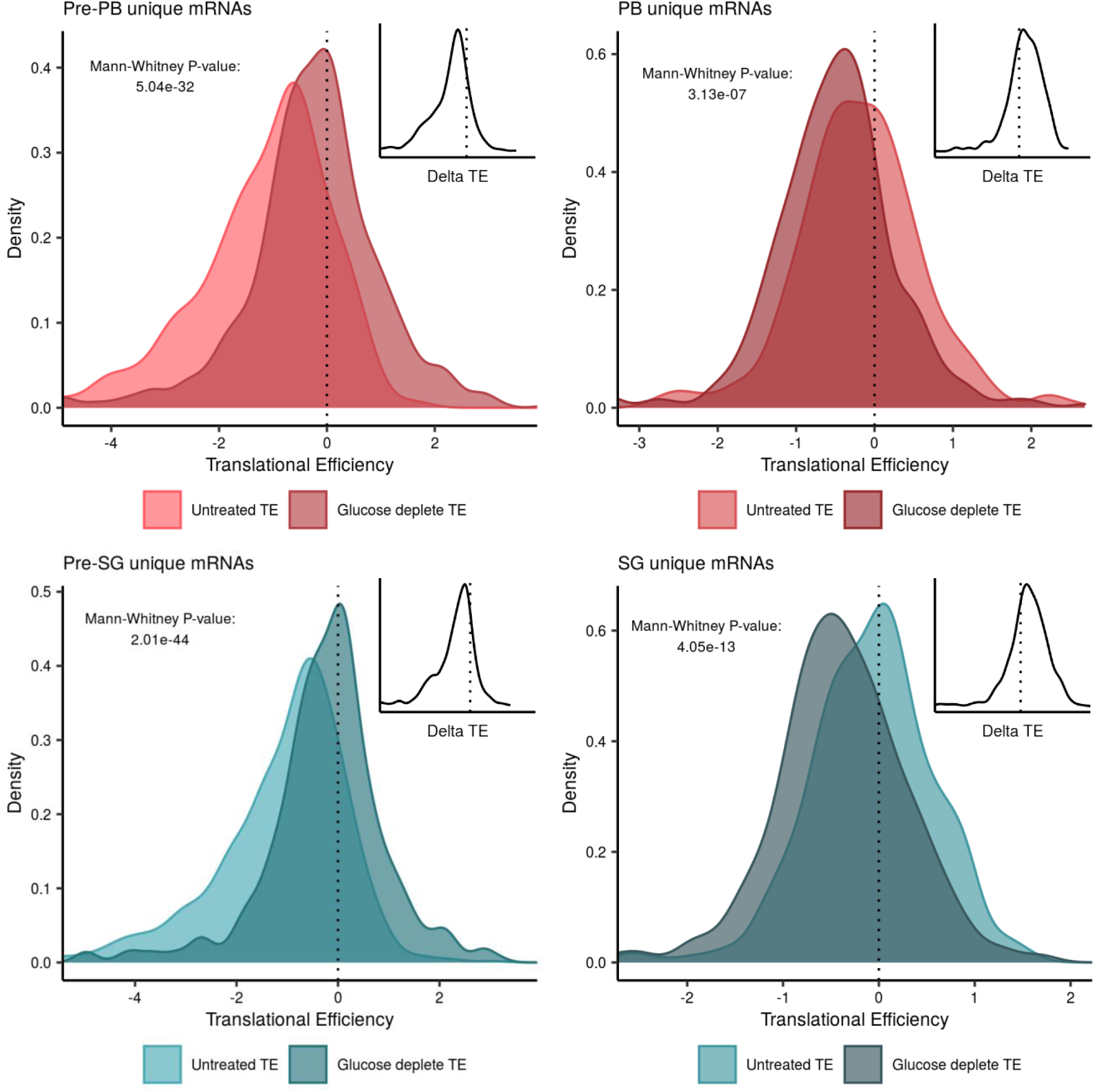
The translational efficiency of RNAs localised to condensates following glucose depletion is reduced. Diagrams are shown comparing TEs determined from unstressed cells or cells following glucose depletion for mRNAs unique to pre-PBs, PBs, pre-SGs and SGs. Inset diagrams show the per mRNA delta between the unstressed and glucose deplete TEs.

Overall as might have been predicted based upon the known functions of Dcp1p and Pbp1p in mRNA decay and processing, mRNAs associated with these factors are on average less well translated, which correlates well with the known predisposition of these condensates towards translationally repressed mRNAs.

### The abundance of mRNAs localised to P-bodies and stress granules is reduced after glucose starvation

Given the decrease in TE observed for mRNAs that localize to PBs and SGs, we determined whether differences in mRNA abundance are also evident using the data from the transcriptomic analysis that we determined as part of our ribosome profiling. We noted that pre-PB and pre-SG associated mRNAs are generally of lower abundance than PB and SG associated mRNAs; a trend that persists after 10 or 60 mins following glucose starvation (Supplementary Fig. 10). This observation is consistent with the ribosome profiling data and suggests that many abundant heavily translated mRNAs are transferred to PBs and SGs after glucose starvation.

To further assess mRNA abundances in the different experiments, we compared the mRNA properties of those that uniquely associate with Dcp1p or Pbp1p, both before and following glucose starvation. This analysis shows an even starker trend. Transcripts uniquely associated with PBs or SGs are significantly more abundant in the total transcriptome than their pre-PB or pre-SG counterparts (Fig. 8A). Although there are no significant differences between the mRNA abundances of the unique condensate transcripts before glucose depletion, in contrast there is a notable difference for those mRNAs unique to PBs and SGs. The mRNAs that uniquely associate with Dcp1p following 10 min glucose show a modest but significant decrease in total cellular abundance compared with unstressed conditions. Additionally, the abundances of the mRNAs that associate with Pbp1p are significantly reduced following 10 and 60 min glucose depletion compared with unstressed conditions (Fig. 8A). Taken together, these data indicate that the mRNAs that relocalize to PBs and SGs following glucose depletion show decreased translational efficiencies and decreased mRNA abundances consistent with gene expression being down-regulated for these mRNAs.

**Fig. 8.**
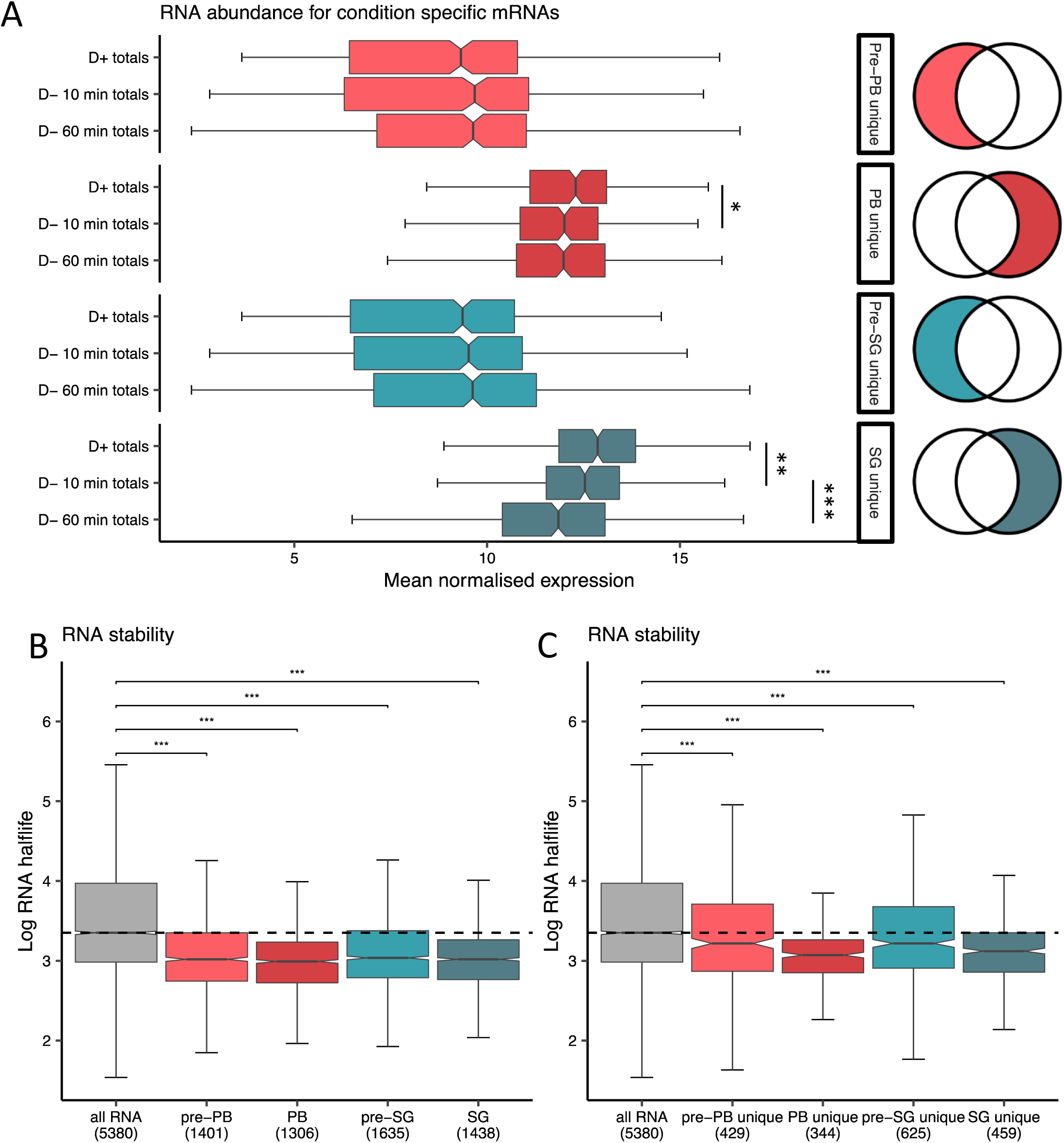
Comparison of abundance and stability for condensate-enriched mRNAs. **A.** Box plots comparing the transcript abundances of mRNAs that are uniquely enriched in pre-PBs, PBs, pre-SGs and SGs under non-stress and following 10 or 60 minutes glucose depletion conditions. **B.** Box plots comparing the RNA stability of mRNAs enriched in pre-PBs, PBs, pre-SGs and SGs with all mRNAs from the transcriptome as a background control. **C.** Box plots comparing the RNA stability of mRNAs that are uniquely enriched in pre-PBs, PBs, pre-SGs and SGs with all mRNAs.

We also considered the stability of mRNA across the different subgroups using previous mRNA half-life measurements obtained from unstressed cells (Neymotin, Athanasiadou et al., 2014). For all of our subgroups, mRNAs with reduced half-lives compared to transcriptome averages are enriched (Fig. 8B). Equally, the half-lives of mRNAs *uniquely* associated with Dcp1p or Pbp1p under unstressed or glucose starvation conditions are also significantly lower than transcriptome averages (Fig. 8C). While these data do not provide direct evidence for degradation occurring in condensates, they do suggest that transcripts localizing to SGs and PBs are typically shorter lived and more readily turned over.

### eIF4A and Ded1-bound mRNAs localise to P-bodies and stress granules

Recent studies have implicated RNA-dependent DEAD-box ATPases in regulating phase separated condensates (Hondele et al 2019, Tauber et al 2020). To evaluate the potential fate of mRNAs interacting with these enzymes, we performed RIP-Seq on Ded1p and eIF4A isolated from untreated or glucose depleted cultures and then cross-compared mRNAs to the pre-PB, pre-SG, PB and SG mRNA pools. Under untreated conditions, the mRNAs that Ded1p binds have strong enrichment scores in pre-PB, PB, pre-SG and SG pools. In comparison, the mRNAs bound to Ded1p following glucose depletion are less likely to be present in these condensates consistent with Ded1 playing a role in modulating the localization of mRNAs to phase separated condensates. In contrast, eIF4A-bound mRNAs from either unstressed or glucose deplete conditions are enriched across all of the condensate-associated mRNA pools (Fig. 9). This could indicate that in contrast to higher eukaryotes (Tauber et al 2020), yeast Ded1 plays a main role in modulating RNA condensation, whilst eIF4A may play a constitutive role in mRNA entry to condensates regardless of cell stress. It should be noted however, that the cellular concentrations of eIF4A are approximately six-fold higher than Ded1p (Kulak, Pichler et al., 2014) meaning that most mRNAs are more likely to be eIF4A-bound irrespective of nutritional conditions. Any role for Ded1p and eIF4A could be either direct via effects on the RNA structure or indirect via effects on mRNA translation. Indeed, both eIF4A and Ded1p may play roles in the rapid inhibition of translation that occurs after glucose depletion (Castelli, Lui et al., 2011, Janapala, Preiss et al., 2019).

**Fig. 9.**
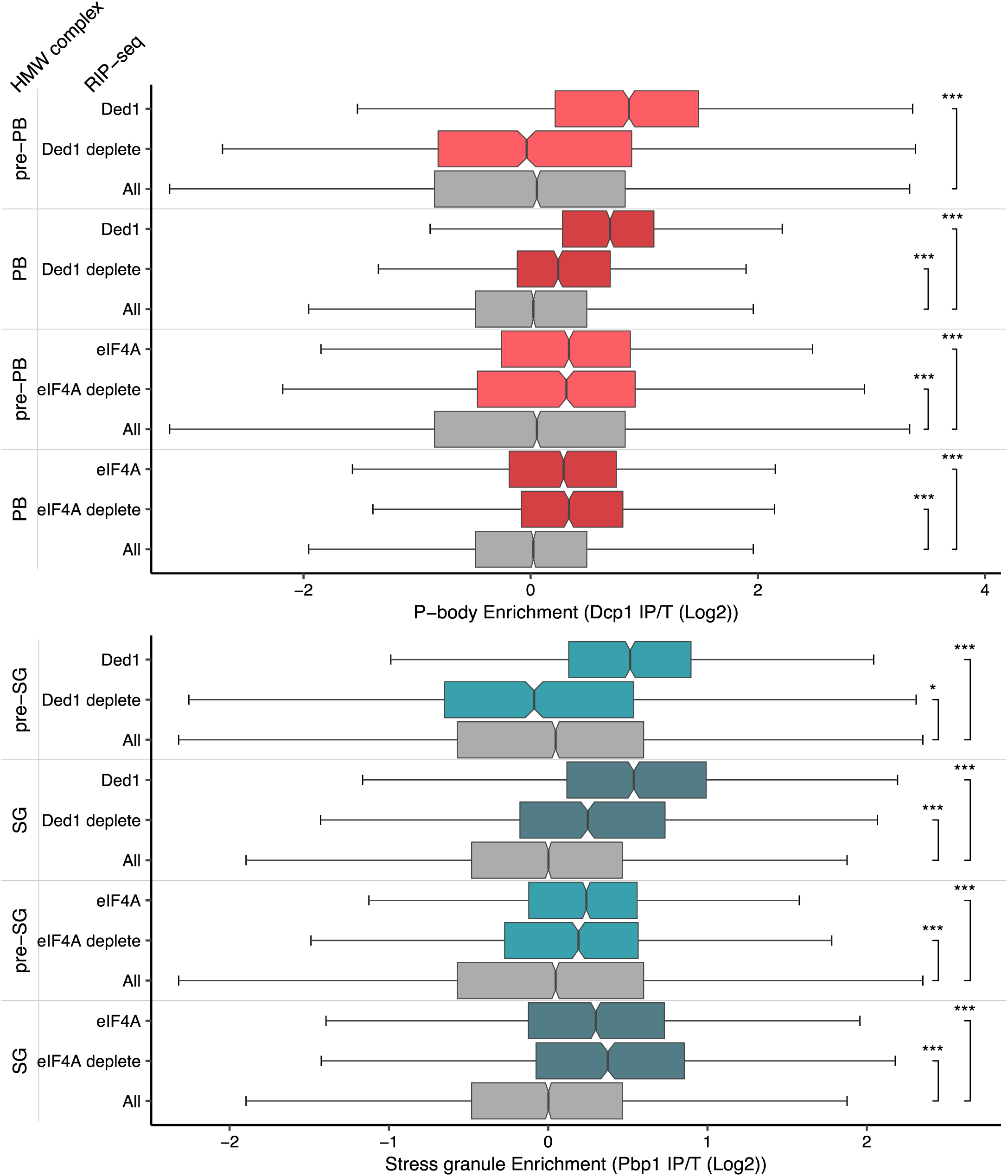
Ded1 and eIF4A associated mRNAs are enriched in PBs and SGs. Boxplots show the enrichment of Ded1p and eIF4A mRNA targets under unstressed and glucose depletion conditions in pre-PBs, PBs, pre-SGs and SGs.

### Diverse RNA binding proteins may play a role in targeting mRNAs to condensates

Previous studies have implicated numerous RNA binding proteins (RBPs) in the formation of PBs and SGs (Jain et al., 2016, Wang et al., 2018). We also identified several known RBPs in PBs and SGs (Fig. 4C and D). This is not entirely surprising given that RNA binding proteins frequently also contain intrinsically disordered domains or regions, and proteins in our purified PBs and SGs are enriched for these properties. To further assess the specificity of RBP involvement in condensate formation, we cross-compared the mRNA sets present in pre-PBs, pre-SGs, PBs and SGs with mRNAs shown to co-immunoprecipitate with various RBPs. For this analysis we used our own RIP-Seq data along with data from a previous comprehensive study by Hogan and colleagues that identified RBP targets (Castelli, Talavera et al., 2015, Costello, Castelli et al., 2015, Costello, Kershaw et al., 2017, Hogan, Riordan et al., 2008, Kershaw, Costello et al., 2015a, Kershaw, Costello et al., 2015b). Strikingly, we noted that the RBPs partitioned into two broad classes; those where the mRNA interactors are enriched across condensates, and those where the mRNA interactors are depleted (Fig. 10). More specifically, this analysis revealed that the mRNA targets of several RBPs including eIF4G1, eIF4G2, Eap1, Sro9, Caf20, Puf3, Vts1, Nop56 and Puf5 localise across pre-PBs, pre-SGs, PBs and SGs (Fig. 10). Scp160 and Puf1 mRNA targets are specifically found in PBs and SGs, Bfr1 mRNA targets are in SGs and Nab2 mRNA targets are in PBs, pre-SGs and SGs. It should be emphasized that it is unclear from these data whether RBPs continue to bind their RNA targets once they are localised to liquid phase separated condensates, but it is consistent with a key role for diverse RBPs in mRNA-condensate localization. Moreover, we noted that some of the proteins themselves were also detected in our proteomics datasets (e.g. Scp160p, eIF4G, Sro9p, Ded1p, Puf3p, Bfr1p, Pab1p, Pub1p, Gbp2, Mrn1p).

**Fig. 10.**
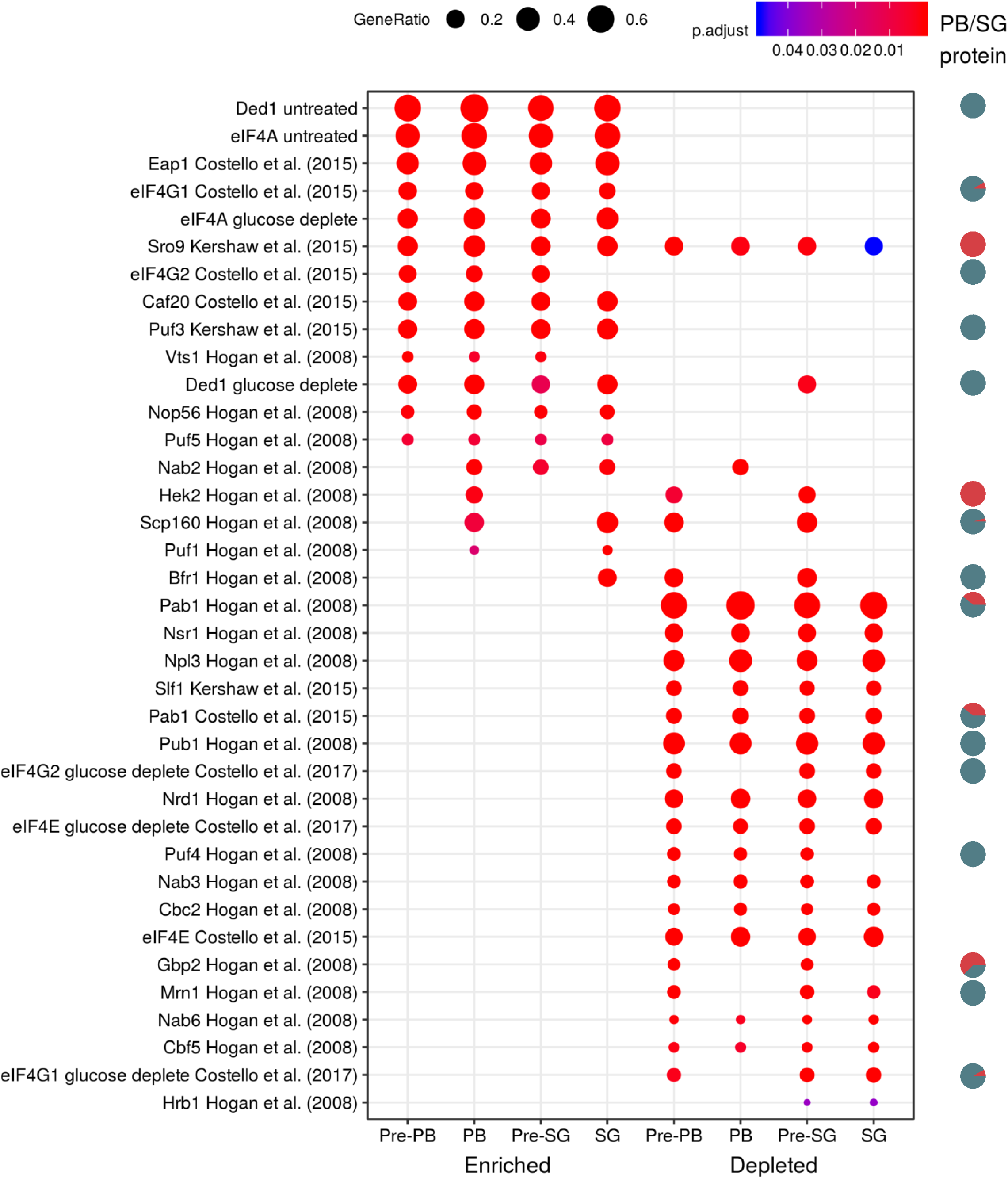
Enrichment of RNA binding protein mRNA targets in PBs and SGs. Comparison of the enrichment or depletion for mRNAs present in pre-PBs, pre-SGs, PBs and SGs with mRNAs previously shown to co-immunoprecipitate with specific RBPs.

As noted above, the mRNA interactors for many RBPs can also be under-enriched in the condensates; for example, those from Pab1p, Nsr1p, Npl3p, Slf1p, Pub1p, Nrd3, eIF4E, Puf4, Nab3 Cbc2p, Nab6p and Cbf5p (Fig. 10). In some cases this seems somewhat surprising since the RBPs themselves apparently localize to PBs and SGs (e.g. Pab1p). However, several are very general RNA binders, interacting with the majority of actively translated transcripts and so any limited RNA subset generated from an independent experiment will appear to be ‘under-enriched’. Equally, many may also act to prevent their mRNA targets from localizing to condensates.

Relatively few RIP-Seq datasets are available from cells grown under glucose depletion conditions. We have previously identified eIF4G1, eIF4G2 and eIF4E mRNA targets following 10 min glucose depletion (Costello et al., 2017). eIF4E targets are under-enriched in pre-PBs, pre-SGs, PBs and SGs regardless of nutritional conditions presumably because eIF4E associated mRNAs are actively translated. Interestingly, whilst eIF4G1 and eIF4G2 targets in unstressed cells tend to localise to pre-PBs, pre-SGs, PBs and SGs, their targets (along with those of eIF4E) after glucose depletion do not and are under-enriched. This result highlights the importance of eIF4G in the dramatic translational reorganisation that occurs after glucose depletion and is in keeping with the rapid alterations in eIF4G interaction with eIF4A and eIF3 coincident with the translation inhibition (Castelli et al., 2011, Janapala et al., 2019).

## CONCLUSIONS

This study presents a first systematic and integrated analysis of the transcriptomes and proteomes of PBs and SGs induced by glucose depletion, as a common stress condition. We provide evidence for the existence of pre-PB and pre-SG ‘seeds’ which after glucose depletion serve as a focus for the condensation of proteins that are remodelled to form mature PBs and SGs, respectively. Our analysis provides support for recent models for RNA / protein condensation, including key physical characteristics of RNA and protein that favour condensation, the role of RNA remodelling and translation control, and a reliance upon a specific cohort of RNA binding proteins. The data also highlight a significant level of interaction and overlap between the constituents found in PBs and SGs. This supports models where common features impact upon both the formation and interaction between these different condensates.

Our quantitative proteomic analysis represents a novel approach to the identification of protein members of hard-to-purify condensates, which we contend offers a more nuanced view on the membership of these phase-separated granules. Our approach considers the quantitative signal associated with proteins in multiple stages of a purification protocol, so that confounding signal arising from fractions other than the key immunoprecipitation itself are also captured and factored into the clustering. This supports the definition of an elution profile across multiple fractions (Fig. 2) that we argue better represents the true subcellular body; much like the LOPIT approach on which it is based (Geladaki, Kocevar Britovsek et al., 2019). In turn, the downstream fuzzy clustering has supported the identification of a previously unrecognized segregation pattern for associated RNA binding proteins. Whilst overlaps between PB and SG components have been noted before in mammalian cells, our approach has now enabled us to quantify the relative contribution of overlapping proteins to either PBs or SGs.

Similar to our proteome analysis, we found that the mRNAs present in condensates appear more dependent on general physical characteristics than the identity of their parent gene, although we do not rule out that particular mRNAs may favour certain condensate localization. The mRNAs that localize to pre-PBs, pre-SGs, PBs and SGs are generally longer and more structured than transcriptome averages. However, a dramatic shift is observed in the properties of the mRNAs that localize to PBs and SGs following glucose depletion compared with their pre-PB and pre-SG counterparts, being longer and less structured. Furthermore, the mRNAs that uniquely localise to PBs and SGs are less translationally competent, with lower TE values in glucose depleted conditions compared to untreated. The converse is true for transcripts only found in the untreated pre-bodies, whose TE values increase following glucose depletion. Similar trends are observed with decreased mRNA abundances, consistent with gene expression being down-regulated for these mRNAs. Whilst this is consistent with PBs and SGs playing a key regulatory role in moderating gene expression to respond to changing growth conditions, our data also highlight that the functional distinctions between these different condensates remains unclear.

The model shown in Fig. 11 presents an analysis of the network of predicted protein interactions (PPI) across all four granule types (pre-PBs, Pre-SGs, PBs, SGs). Our proteomics analysis suggests that pre-PBs and pre-SGs are entirely independent protein complexes that share no proteins but pre-exist in cells. Relatively few PPIs have previously been identified between the protein components of pre-PBs and pre-SGs suggesting that our analysis has identified a number of novel PPIs for these proteins. Following stress, the pre-PBs and pre-SGs undergo significant remodelling forming PBs and SGs, which contain granule specific proteins as well as overlapping proteins. The formation of compositionally distinct condensates from independent seeds is suggestive of distinct pathways of formation. Although, the fact that there are numerous overlapping protein and RNA components suggests that significant interaction occurs between PBs and SGs, and this interaction could occur at any point during the formation of the mature condensates. Such interaction between the pathways of PB and SG formation could in turn act to facilitate a tuneable response to stress, where protein components are distributed between different condensates rather than being specifically localized to PBs or SGs in a binary manner. A key question is how specificity in PB and SG formation arises, but may for example, be driven by the unique RNA species that populate these granules.

**Fig. 11.**
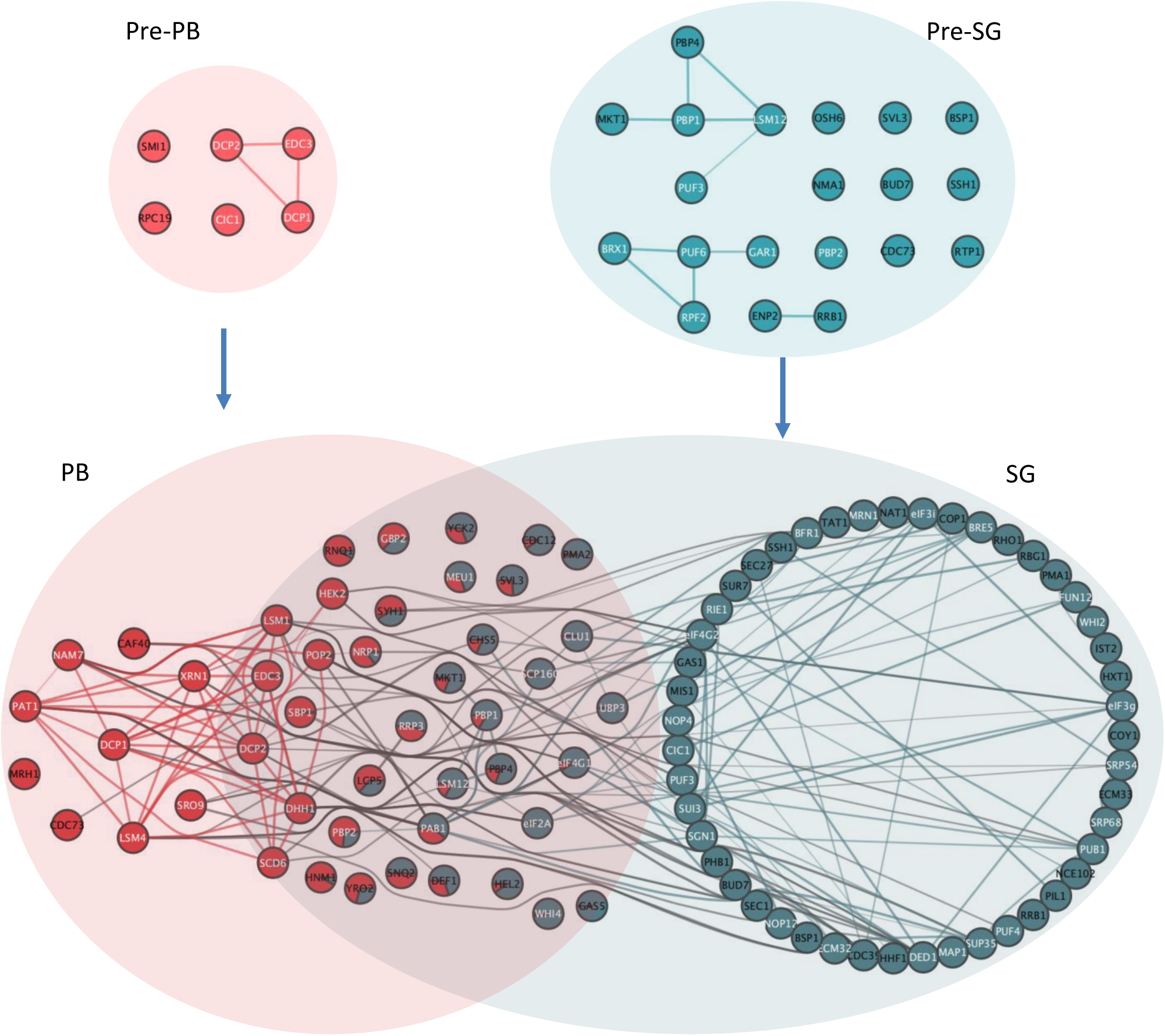
Model of condensation of biomolecules into PBs and SGs. Overlaps between the networks of predicted protein interactions (PPI) for pre-PBs, pre-SGs, PBs and SGs are shown. Implicit PPIs between PB and SG members are contained within each coloured ellipsoids. Each node represents a protein identified as a member of an HMW complex cluster and previously identified protein:protein interactions are indicated by lines between the nodes. Proteins overlapping granules are represented by pie charts depicting cluster membership as in Fig 3E. Interactions between nodes both assigned uniquely or predominantly to the same granule type are coloured the same as the granule. Interactions between nodes uniquely or predominantly of different granules are coloured grey. Proteins labelled in white are those that have an RNA binding GO annotation.

The protein interaction network for our SG and PB components is also consistent with recent studies where graph theory approaches have been used to provide a mechanistic understanding of how multivalent RNA binding proteins promote liquid-liquid phase separation (LLPS) (Guillen-Boixet, Kopach et al., 2020, Sanders, Kedersha et al., 2020, Yang, Mathieu et al., 2020). We find numerous multivalent proteins that may act as hubs to promote PB and SG formation (Fig. 11), highlighting previously established physical PPIs both within and between condensate types. We also find evidence for bridging proteins with double valences (two interaction partners) and capping proteins with single predicted PPIs populating the network of PPIs. Further studies will be required to understand how these multifarious interacting proteins contribute to the specificity of condensate formation, and particularly how PB and SG interactions are bridged by overlapping components.

## MATERIALS AND METHODS

### Yeast strains and growth conditions

*S. cerevisiae* was cultured in SCD complete media (ForMedium, UK) either in the presence or absence of 2% w/v glucose. All growth was performed at 30°C with shaking. Myc-tagged strains were constructed in yeast strain W303-1A *(MAT*a *ura3-52 leu2-3 leu2-112 trp1-1 ade2-1 his3-11 can1-100*) using a PCR-based strategy (Knop, Siegers et al., 1999). Ded1-TAP and eIF4A-TAP strains were obtained from Thermo Scientific Open Biosystems (Waltham, MA, USA). A BY4741 *HIS3* strain (Costello et al., 2015) was used to perform the ribosome profiling.

### Immunoprecipitation of tagged proteins

Immunoprecipitations were performed as previously described (Rowe et al 2014). Briefly, cells were grown to an OD_600_ of 0.6-0.8, before being split and transferred to centrifuge tubes. Cells were collected by centrifugation and resuspended in pre-warmed media (with or without glucose) in pre-warmed conical flasks and incubated for 10 or 60 minutes at 30°C. Cultures were then transferred to chilled centrifuge tubes and crosslinked using 0.8% formaldehyde whilst simultaneously being rapidly chilled by adding frozen media (with or without glucose) to the culture and submerging the centrifuge tube in water ice. Cells were crosslinked for 1 h and then quenched with 0.1 M glycine pH7. Yeast cells were pelleted by centrifugation, washed in 10 ml of Blob100 buffer (20 mM Tris-HCl (pH 8), 100 mM NaCl, 1 mM MgCl_2_, 0.5% (v/v) NP40, 0.5 mM TCEP, EDTA free Protease Inhibitor cocktail tablet (Roche Diagnostics, Indianaplois, IN, USA)). Cells were resuspended in 4 ml of Blob100 buffer plus 80 units/ml RNasin Plus (Promega, Flitchburg, WI, USA) and lysed under liquid nitrogen in a 6870 Freezer Mill (Spex, Metuchan, NJ, USA).

RNA and protein were isolated from the same cultures. In both cases, cell debris was cleared by centrifugation for 10 mins at 1,000g, protein concentration was determined by Bradford assay and then either 2 mg or 4 mg of total protein was used for protein isolation or RNA isolation, respectively. The cleared lysate was centrifuged at 20,000g for 10 mins, washed in 500 μl of Blob100 buffer, and then the spin was repeated. The resulting washed pellet was resuspended in 500 μl of Blob100 buffer.

Immunoprecipitations of protein for mass spectroscopy were performed as follows: 150 ul of anti-c-Myc magnetic beads (Pierce) were washed twice in 500 ul of Blob100 buffer and 150 ug of cleared lysate was applied to the washed myc beads. Bound proteins were eluted using Myc peptide. Prior to RNA isolation, bound RNA was eluted from the beads using Proteinase K (New England Biolabs) at 55^0^C for 15 mins in modified Blob100 buffer (in the presence of 0.5% (w/v) SDS and 1 mM EDTA, but without RNAsin and MgCl_2_). Crosslinks were reversed at 70°C for 40 mins. RNA was isolated using Trizol LS (Invitrogen) as previously described (Costello et al., 2015).

Immunoprecipitation of TAP-tagged proteins was performed as previously described (Costello et al., 2015).

### Mass spectrometry analysis

Protein samples were briefly separated by SDS-PAGE. Each sample was excised in its entirety and dehydrated using acetonitrile followed by vacuum centrifugation. Dried gel pieces were reduced with 10 mM dithiothreitol and alkylated with 55 mM iodoacetamide. Gel pieces were then washed with 25 mM ammonium bicarbonate followed by acetonitrile. This was repeated, and the gel pieces dried by vacuum centrifugation. Samples were digested with trypsin overnight at 37°C.

Digested samples were analysed by LC-MS/MS using an UltiMate^®^ 3000 Rapid Separation LC (RSLC, Dionex Corporation, Sunnyvale, CA) coupled to a QE HF (Thermo Fisher Scientific, Waltham, MA) mass spectrometer. Mobile phase A was 0.1% formic acid in water and mobile phase B was 0.1% formic acid in acetonitrile and the column used was a 75 mm x 250 μm i.d. 1.7 μM CSH C18, analytical column (Waters). A 1ul aliquot of the sample was transferred to a 5ul loop and loaded on to the column at a flow of 300 nl/min for 5 minutes at 5% B. The loop was then taken out of line and the flow was reduced from 300 nl/min to 200 nl/min in 0.5 minute. Peptides were separated using a gradient that went from 5% to 18% B in 63.5 minutes, then from 18% to 27% B in 8 minutes and finally from 27% B to 60% B in 1 minute. The column was washed at 60% B for 3 minutes before re-equilibration to 5% B in 1 minute. At 85 minutes the flow was increased to 300 nl/min until the end of the run at 90min.

Mass spectrometry data was acquired in a data directed manner for 90 minutes in positive mode. Peptides were selected for fragmentation automatically by data dependant analysis on a basis of the top 12 peptides with m/z between 300 to 1750Th and a charge state of 2, 3 or 4 with a dynamic exclusion set at 15 sec. The MS Resolution was set at 120,000 with an AGC target of 3e6 and a maximum fill time set at 20 ms. The MS2 Resolution was set to 30,000, with an AGC target of 2e5, a maximum fill time of 45 ms, isolation window of 1.3Th and a collision energy of 28.

### MS Data analysis

For label-free quantification (LFQ), data were processed using MaxQuant (version 1.6.3.4) (Cox et al., 2014). Raw data were searched against the *S. cerevisiae* W303 protein sequence file (Song, Dickins et al., 2015) obtained from https://downloads.yeastgenome.org/sequence/strains/W303/W303_SGD_2015_JRIU00000000/, and MaxQuant’s reversed decoy dataset and inbuilt set of known contaminants. Default search parameters were used with standard tryptic digestion allowing two missed cleavages and minimum peptide lengths of six. Carbamidomethyl cysteine was specified as a fixed modification. Oxidation of methionine, N-terminal protein acetylation and phosphorylation of serine, threonine and tyrosine were specified as variable modifications. Additionally, “match between runs” was enabled, but limited to occur only between biological replicates. Searches were constrained to 1% FDR at all levels.

An approach was adapted from the hyperLOPIT protocol developed by Lilley and colleagues (Geladaki et al., 2019) for identification of members of HMW complexes, substituting fractions collected from the centrifugation and immunoprecipitation steps for ultracentrifugation. First, the MaxQuant ‘peptides.txt’ output file was processed using the MSnbase R packge (Gatto & Lilley, 2012). Proteins lacking any non-zero measurement in any elution fraction were discarded. Any missing data for the remaining proteins was imputed using QRILC (quantile regression imputation of left-censored data), implemented in the MsnBase package (Gatto & Lilley, 2012). Peptide level quantifications were rolled up to the protein level, as MaxQuant LFQ values, averaged across replicates and sum normalised. Processed data were converted to a Bioconductor *ExpressionSet* class in R (Huber, Carey et al., 2015) and split into untreated and glucose deplete samples for further analysis.

Mfuzz was used to assign proteins to fuzzy clusters, for each of the separate untreated and glucose deplete conditions, in order to infer presence in independent HMW complexes (Kumar & M, 2007). Protein levels across the two sample sets (untreated, glucose deplete) were standardised to a mean value of zero and a standard deviation of one. An optimal fuzzifier *m* was estimated using the Mfuzz *mestimate* function. The number of clusters was determined for each condition by increasing the number until the two tagged proteins, Dcp1 and Pbp1, were assigned to different clusters. Each protein is assigned a membership value for each cluster, and those proteins whose highest score was to one of the two clusters containing the marker proteins were assigned to that HMW complex. Proteins were defined as members of *both* PBs and SGs if the two highest cluster membership scores were for the two marker protein clusters and the lower membership score was at least 0.001. A t-SNE plot was used to visualise the data with cluster membership used for colouring data points (Krijthe, 2015, van der Maaten, 2014).

For comparison with our clustering approach, a classic IP enrichment analysis was performed using Perseus (version 1.6.2.2) (Tyanova, Temu et al., 2016). The MaxQuant “proteinGroups.txt” file was loaded into Perseus with LFQ columns relating to the tagged IP and untagged IP selected as main columns. Reverse hits, contaminants and “only identified by site” were filtered out. Next, log2 values were taken and missing values imputed from a normal distribution shrunk by a factor of “0.3” (width) and shifted down by “1.8” (down shift) standard deviations. Finally, interactors were determined by a two sample T-test using s0 = 2 and FDR cut-off of 0.01. The data from these analyses are presented in Supplementary Tables 3 and 4.

Gene ontology over-representation was calculated using the R Bioconductor package *clusterProfiler* (Yu, Wang et al., 2012). The probability of containing a long IDR was calculated using SLIDER (Peng, Mizianty et al., 2014). The proportion of disordered amino acids was calculated by DISOPRED3 (Jones & Cozzetto, 2015). For both analyses, all proteins detected by mass spectrometry were used as the background set for Mann-Whitney statistical significance. Protein-protein interaction networks for proteins identified in granules were created in Cytoscape (Shannon, Markiel et al., 2003) using data from STRING (Szklarczyk, Gable et al., 2019) with only experimental interactions with a confidence of greater than 0.4 drawn. RNA binding annotations highlighted on networks are defined as proteins annotated with the Molecular Function Gene Ontology term “RNA binding” or any daughter term. Protein properties were calculated using Composition Profiler (Vacic, Uversky et al., 2007).

### Next generation sequencing

Total RNA was submitted to the University of Manchester Genomic Technologies Core Facility (GTCF). Quality and integrity of the RNA samples were assessed using a 2200 TapeStation (Agilent Technologies) and then libraries generated using the TruSeq® Stranded mRNA assay (Illumina, Inc.) according to the manufacturer’s protocol. Briefly, rRNA depleted total RNA (0.1-4ug) was used as input material which was fragmented using divalent cations under elevated temperature and then reverse transcribed into first strand cDNA using random primers. Second strand cDNA was then synthesised using DNA Polymerase I and RNase H. Following a single ‘A’ base addition, adapters were ligated to the cDNA fragments, and the products then purified and enriched by PCR to create the final cDNA library. Adapter indices were used to multiplex libraries, which were pooled prior to cluster generation using a cBot instrument. The loaded flow-cell was then paired-end sequenced (76 + 76 cycles, plus indices) on an Illumina HiSeq4000 instrument. Finally, the output data was demultiplexed (allowing one mismatch) and BCL-to-Fastq conversion performed using Illumina’s bcl2fastq software, version 2.17.1.14

As the *S. cerevisiae* genome contains few UTR annotations, we built an annotation set based on TIF-Seq data from (Pelechano, Wei et al., 2013). For each ORF in the Ensembl version R64-1-1 annotation, the UTRs with the highest read support on YPD from (Pelechano et al., 2013) was selected and used as our UTR annotation. Fastq files of all reads were mapped to the *S. cerevisiae* genome using the splice aware STAR aligner (version 2.5.3) (Dobin, Davis et al., 2013) with our modified UTR annotation. Uniquely mapping reads were retained and stored in BAM format by samtools (version 1.9) (Li, Handsaker et al., 2009). Mapped counts per gene were calculated by htseq-count (version 0.10) (Anders, Pyl et al., 2015). Raw counts were processed using DESeq2 (Love, Huber et al., 2014) with enrichments calculated in the IP samples with respect to the matched condition total. When performing pairwise analysis on the RIP-Seq data, data was reduced to the maximum number of mRNAs that were sequenced in both samples. For example, when comparing unstressed Pbp1p (6718) vs Pbp1p glucose deplete (6839) only the 6718 present in both were used for the analysis. Enriched gene lists were selected with an adjusted p-value of less than 0.01.

Gene ontology over-representation was calculated using clusterProfiler, based on GO slim mappings obtained from SGD (https://downloads.yeastgenome.org/curation/literature/). UTR lengths (where annotated) were taken from the same modified annotation used for RNA mapping. Published datasets were used for PolyA tail lengths (Subtelny, Eichhorn et al., 2014), RNA structure gini index (Rouskin et al., 2014) and RNA stability (Neymotin et al., 2014). Theoretical translation efficiency was calculated using a tRNA adaptation index calculation termed the classical translational efficiency (cTE), which uses codon optimality based on tRNA gene copy numbers and codon usage in a subset of highly expressed genes (dos Reis, Savva et al., 2004, Pechmann & Frydman, 2013). VST transformed RNA abundance was extracted from the DESeq2 model and the median taken from the appropriate total samples for each condition. Background lists were either all mRNA or all RNA as labelled. Enrichment and depletion of HMW complex enriched RNAs in previous RIP-Seq experiments was calculated using the clusterProfiler ‘enricher’ function for custom lists.

### Ribosome profiling and polysome analysis

Polysome analysis was performed by taking a small sample of the extracts made for protein and RNA isolation that was cleared by centrifugation at 1,000g for 10 mins. The RNA concentration of this cleared lysate was determined using a Nanodrop and 2.5 A_260_ units were loaded onto a sucrose gradient. 15–50% sucrose gradients were poured as previously described (Taylor, Campbell et al., 2010) and the A_256_ was continuously read to produce the trace. Fractions were taken every 30 s and protein was precipitated using an equal volume of 20% TCA.

Ribosome profiling (McGlincy & Ingolia, 2017) was performed using the TruSeq Ribo Profile (Yeast) kit (Illumina, San Diego, USA). A BY4741 *HIS3* strain (Costello et al., 2015) was grown to an OD_600_ 0.6-0.8 and either untreated or glucose starved for 10 minutes. The manufacturer’s protocol was followed with the exception of the purification of the 80S monosome where sucrose density centrifugation was used rather than a MicoSpin column. Fastq files of all reads were mapped to the *S. cerevisiae* genome using the splice aware STAR aligner (version 2.5.3) with our modified UTR annotation. Uniquely mapping reads were retained and stored in BAM format by samtools (version 1.9). Mapped counts per gene were calculated by htseq-count (version 0.10). Raw counts were processed using DESeq2, with TE calculated as the Log_2_ fold change between ribosome footprint samples with respect to the matched condition total.

### Western blot analysis

IP samples were resolved by SDS–PAGE, electroblotted onto nitrocellulose membrane and probed using the relevant primary antibody. Bound antibody was visualized using WesternSure Chemiluminescent Reagents (LI-COR).

## Data availability

The datasets produced in this study are available in the following databases:

- Granule RIP-Seq data: ArrayExpress E-MTAB9096 (https://www.ebi.ac.uk/arrayexpress/experiments/E-MTAB-9096)
- Ded1/eIF4A RIP-Seq data: ArrayExpress E-MTAB-9095 (https://www.ebi.ac.uk/arrayexpress/experiments/E-MTAB-9095)
- Ribo-Seq data: ArrayExpress E-MTAB-9094 (https://www.ebi.ac.uk/arrayexpress/experiments/E-MTAB-9094)
- Protein mass spectrometry data: PRIDE PXD018762 (https://www.ebi.ac.uk/pride/archive/projects/PXD018762)

## Acknowledgements

We would like to thank the Genomic Technologies Core Facility and the Bio-MS Research Core Facility in Faculty of Biology, Medicine and Health at The University of Manchester for technical help and useful discussions. This work was supported by grant BB/P005594/1 from the UK Biotechnology and Biological Sciences Research Council (www.bbsrc.ac.uk). JL was funded by BBSRC project grant BB/K005979/1 and MDJ by BBSRC project grant BB/N014049/1. CB was funded by a WT PhD studentship 210002/Z/17/Z. Additional support for core research facilities was provided by Wellcome Trust Institutional Strategic Support Fund awards (www.wellcome.ac.uk). The funders had no role in study design, data collection and analysis, decision to publish, or preparation of the manuscript.

## Authors’ contributions

Conceived and designed the experiments: CMG, MPA, SJH, MGN and CJK. Performed the experiments: CJK, JL, CPB, MDJ. Analysed the data: MGN, CJK CMG, MPA and SJH. Wrote the paper: CJK, CMG, MPA, SJH and MGN. All authors were involved in intellectual aspects of the study, and they edited and approved the final manuscript.

---

**Supplementary Fig. 1.**
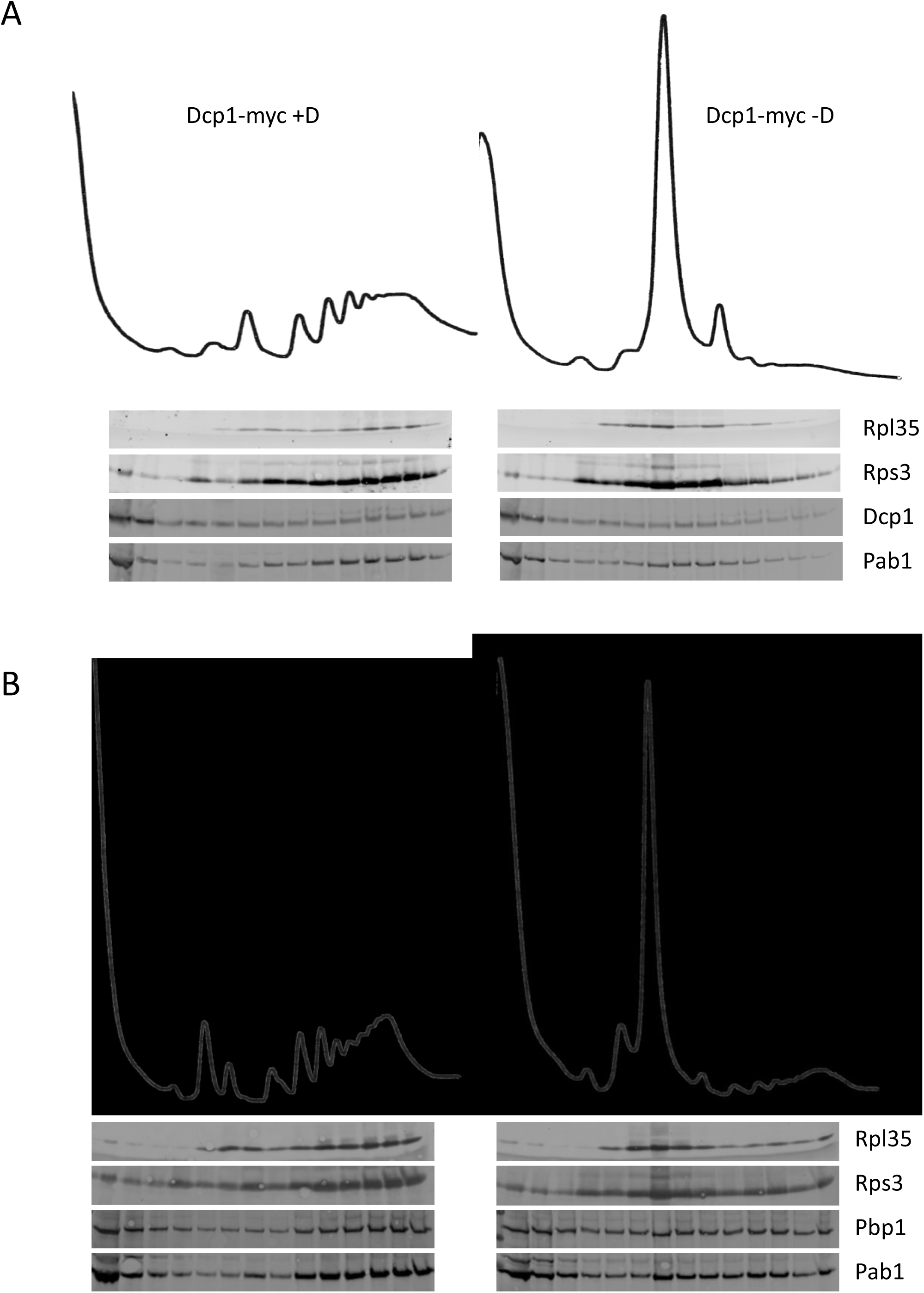
Poly-ribosome analysis of tagged strains under unstressed and glucose depletion conditions. **A, B.** Poly-ribosome profiles for Dcp1p-myc and Pbp1p-myc strains from untreated (+D) or glucose depletion (-D) conditions. Western blotting is shown for fractions isolated from polyribosome gradients probed against the indicated proteins. Glucose depletion was imposed for 10 or 60 minutes for Dcp1p-myc and Pbp1p-myc strains, respectively.

**Supplementary Fig. 2.**
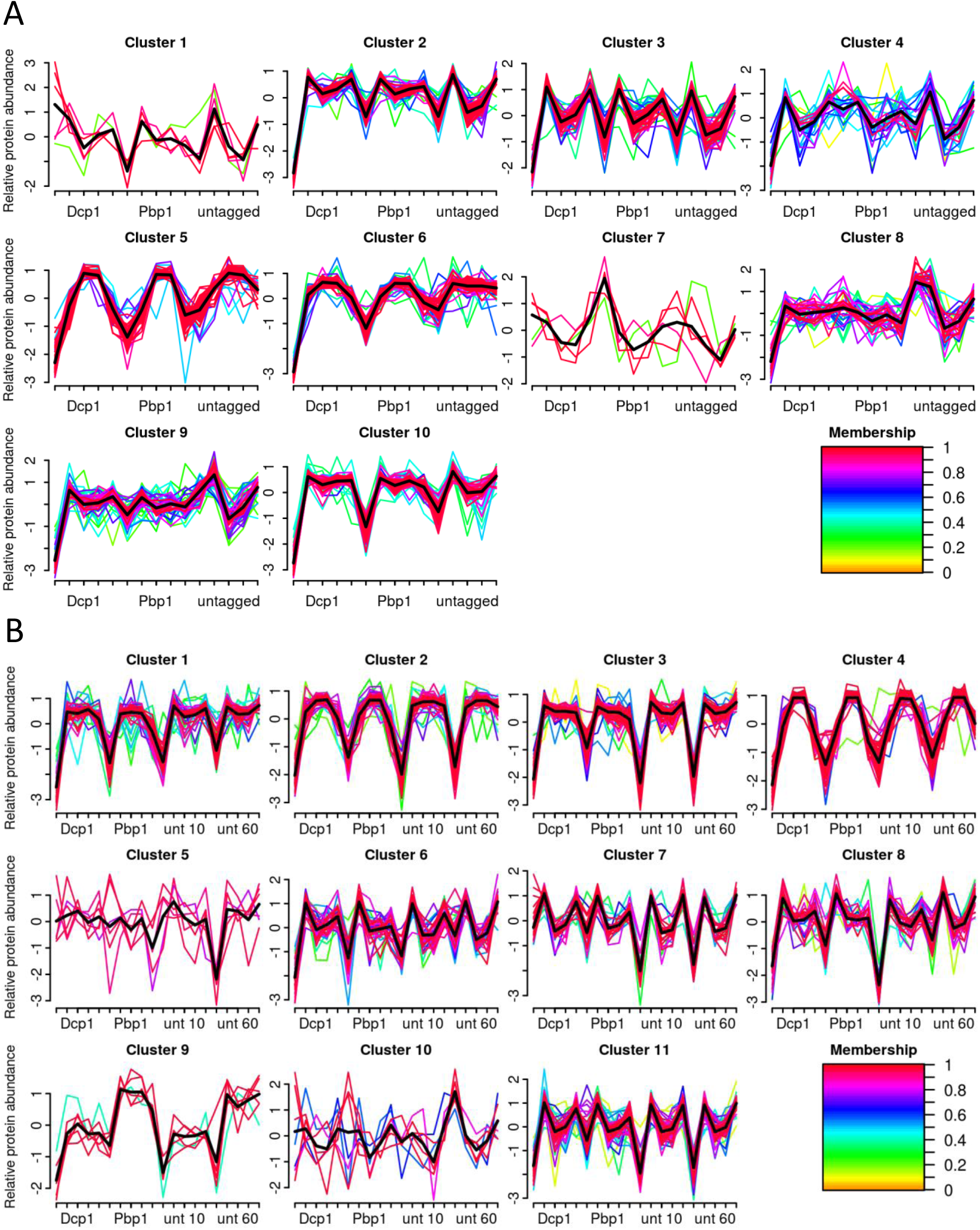
Cluster analysis used to identify condensate components. **A, B.** Clustering of proteomic data resulted in 10 clusters for untreated and 11 clusters for glucose deplete samples. Each x-axis label represents a block of five normalised protein signals from IP, pellet, supernatant, total and unbound fractions, respectively, with 15 fractions in unstressed and 20 fractions in glucose deplete conditions. Known PB and SG components were used to designate clusters representing pre-PBs (Cluster 1), PBs (Cluster 7), pre-SGs (Cluster 4), and SGs (Cluster 11). Lines are coloured by how well each protein correlates with the cluster and a black line represents the average of all data within the cluster.

**Supplementary Fig. 3.**
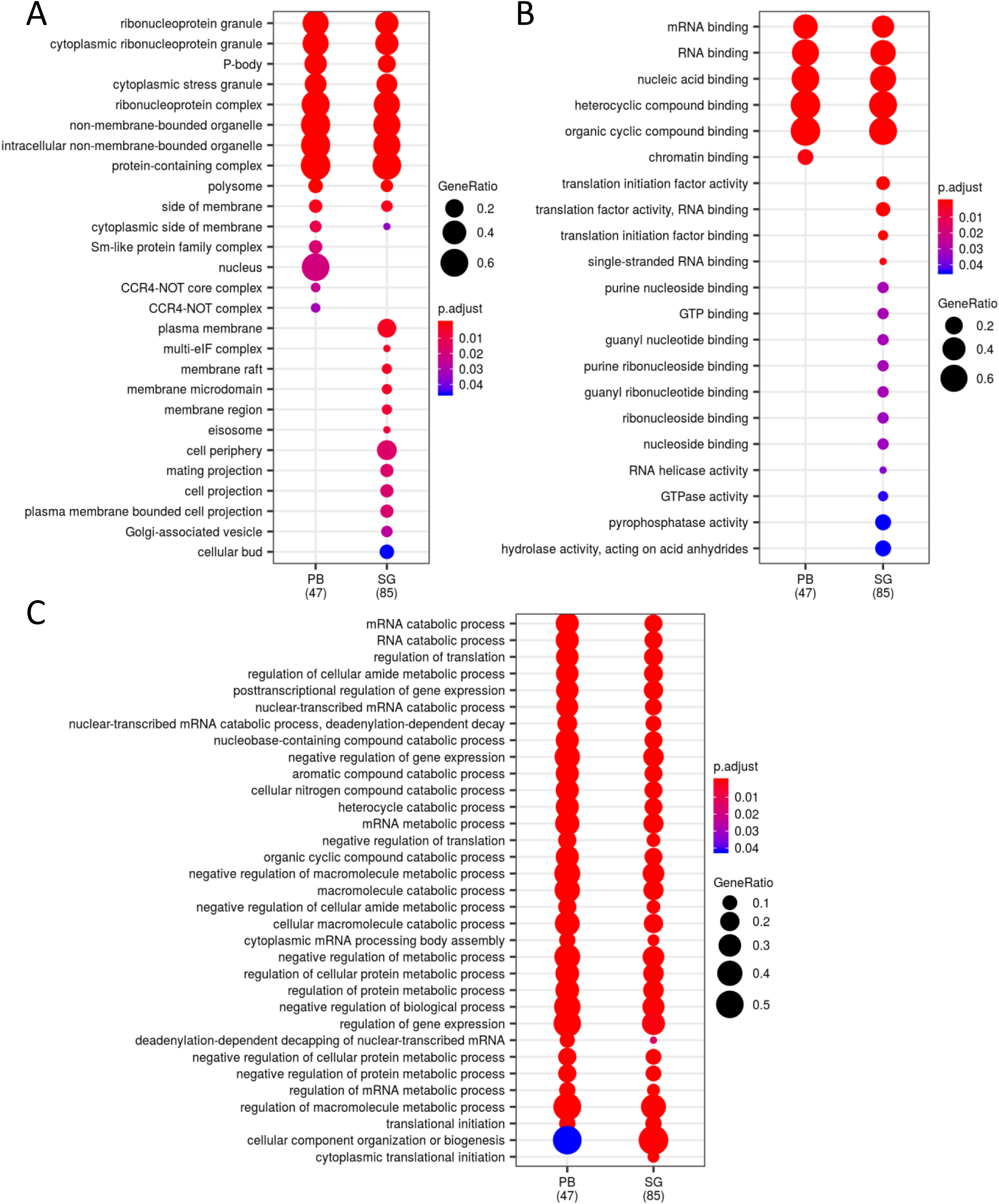
Functional categorisation of proteins present in PBs and SGs following glucose depletion.

**Supplementary Fig. 4.**
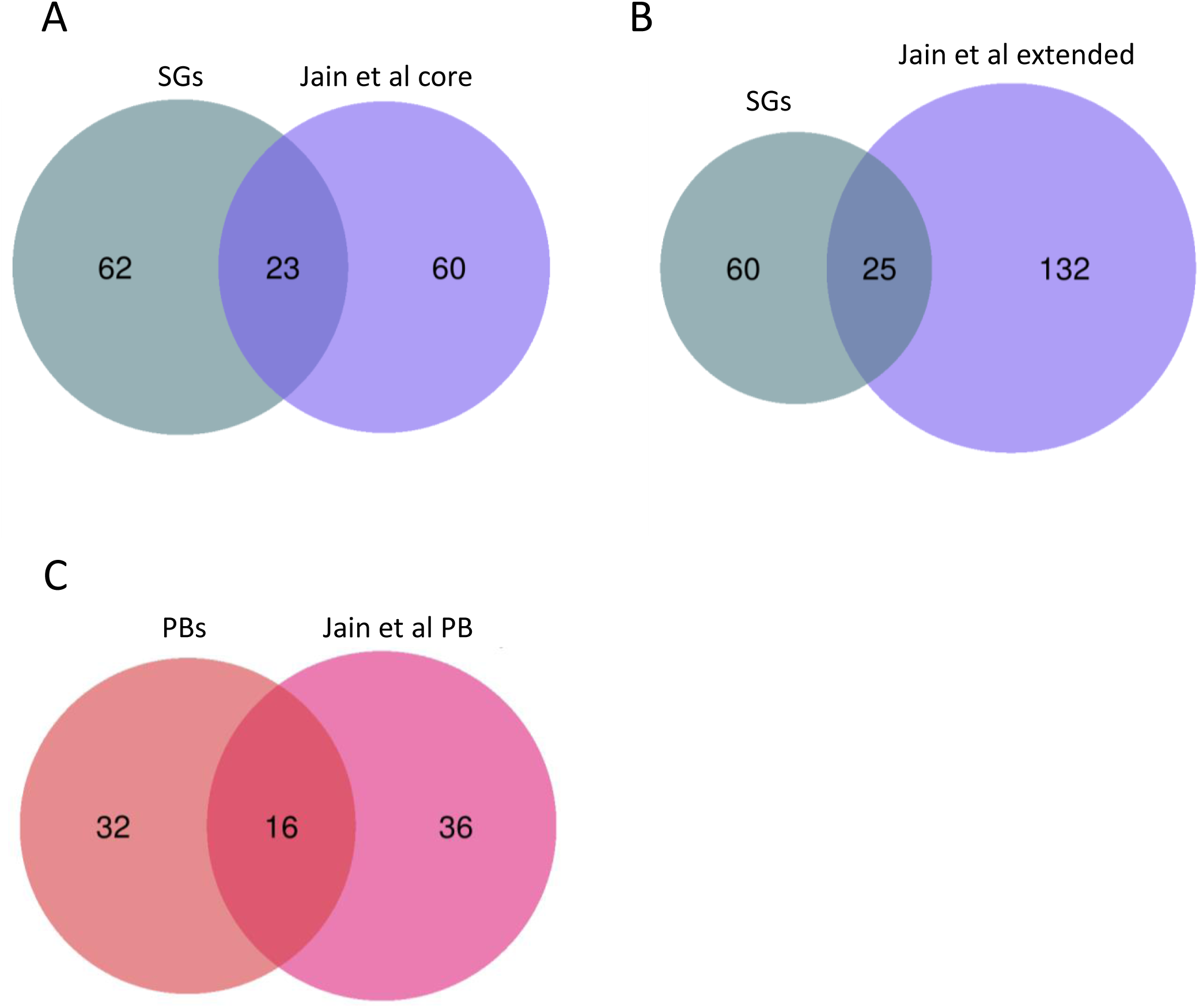
Comparison of glucose-depletion induced SG and PB-associated proteins with other published condensate proteomes. **A, B.** Euler diagrams comparing the protein components in our glucose-depletion SGs with previously published ‘core’ (**A**) and ‘extended’ SG (**B**) proteomes (Jain et al., 2016) induced by sodium azide stress. **C.** Euler diagram comparing the protein components in our glucose-depletion PBs with a literature-mining based analysis of PB associated proteins (Jain et al., 2016).

**Supplementary Fig. 5.**
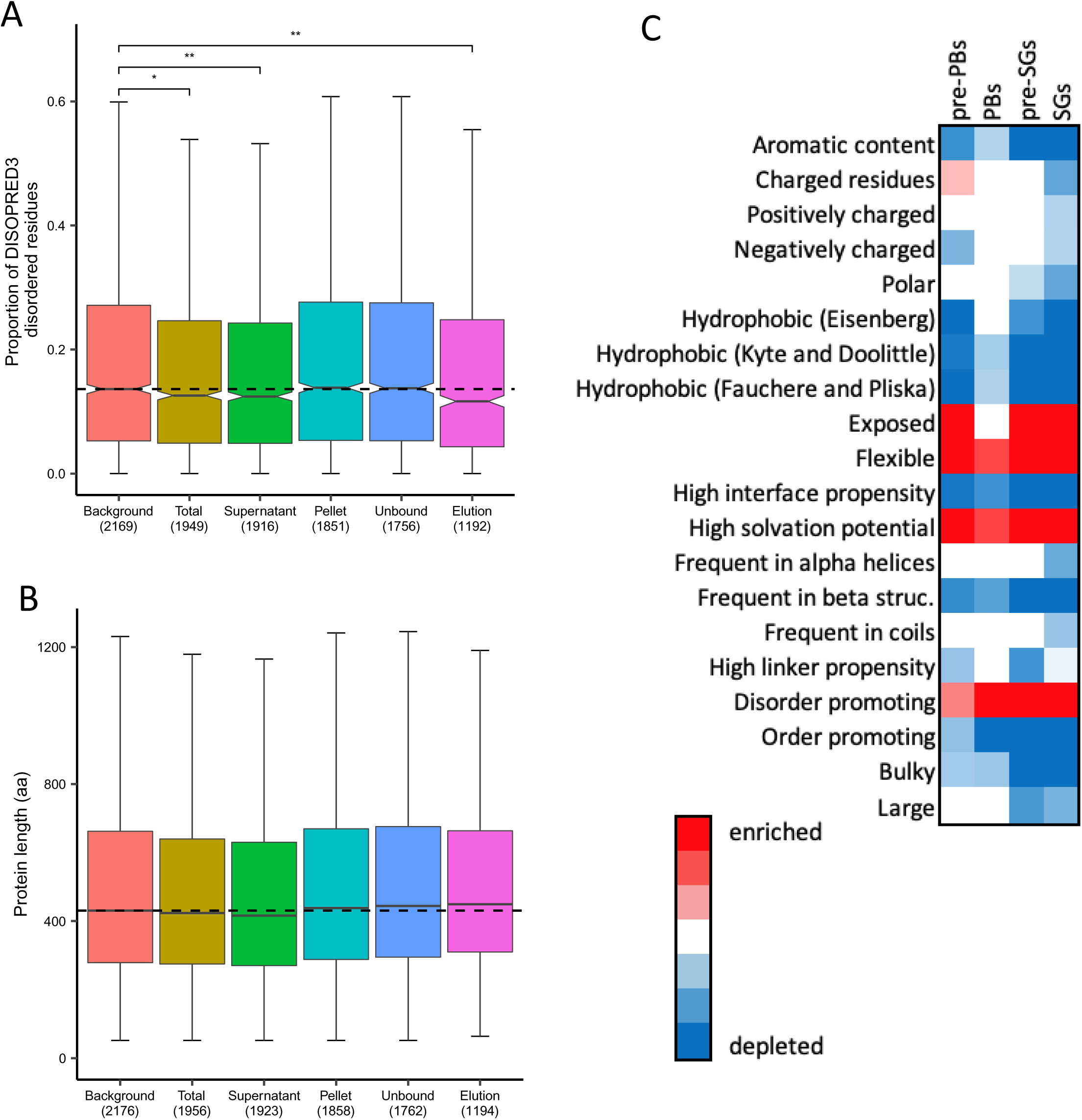
Comparison of protein disorder, protein length and amino acid biophysical properties in condensate proteomes. **A.** Box-plots comparing the probability of condensate proteins containing disordered (as determined by DISOPRED3) residues for the various fractions isolated during the condensate preparation process. The background dataset used was all proteins present in the mass spectroscopy analysis. **B.** Box-plots comparing the protein lengths for the various fractions isolated during the condensate preparation process. **C.** Heat map showing confidence (*p*-values) for the biophysical properties of those amino acids that are enriched (red) and those that are depleted (blue) in PBs and SGs isolated from unstressed conditions or following glucose depletion.

**Supplementary Fig. 6.**
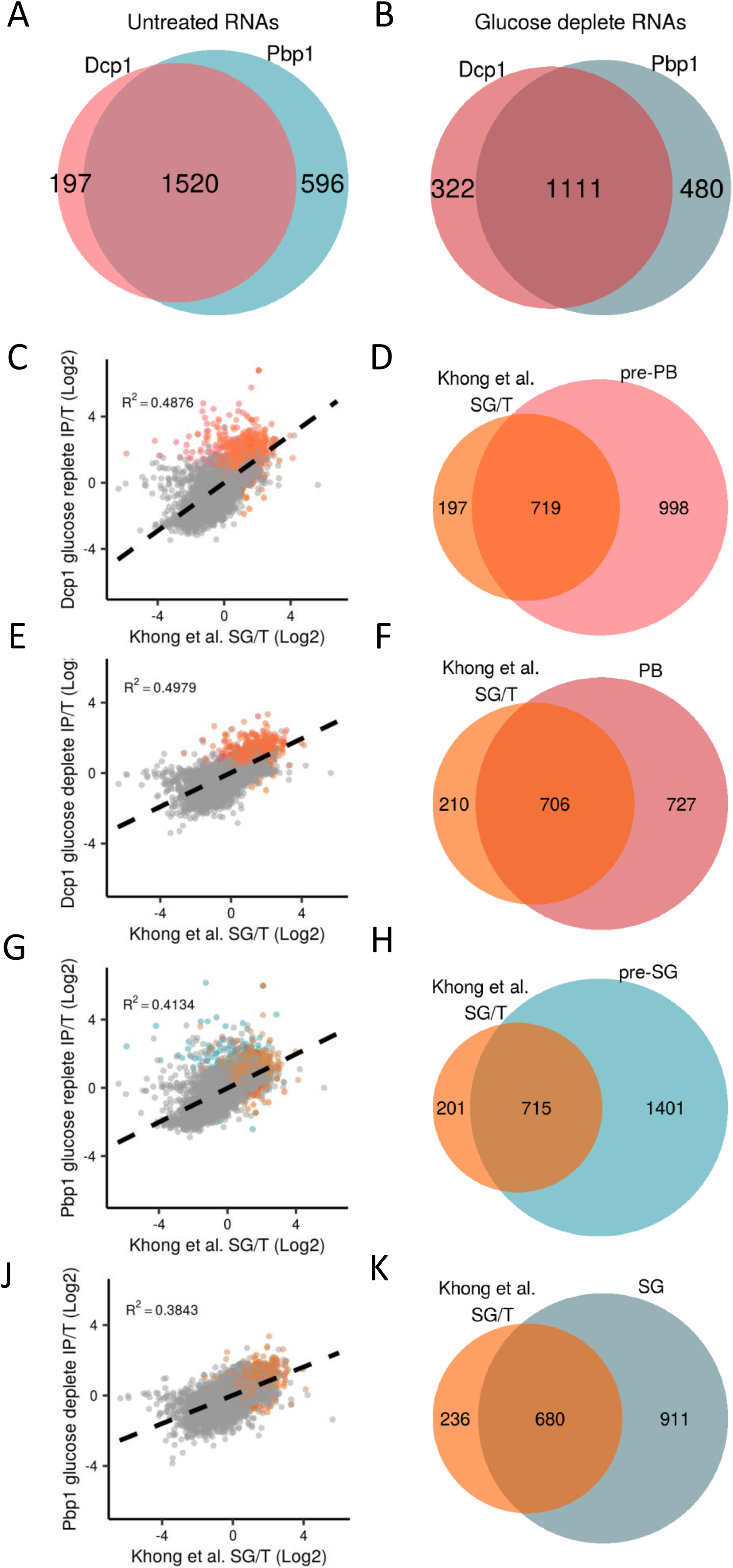
Comparison of PB and SG transcriptomes. **A.** Euler diagram comparing mRNA components of pre-PBs and pre-SGs **B.** Euler diagram comparing mRNA components of PBs and SGs **C-K.** XY scatterplots and Euler diagrams are shown comparing Log_2_ fold enrichment of IP/T for the mRNAs in our HMW complexes with mRNAs identified in sodium azide induced PBs (Khong et al., 2017).

**Supplementary Fig. 7.**
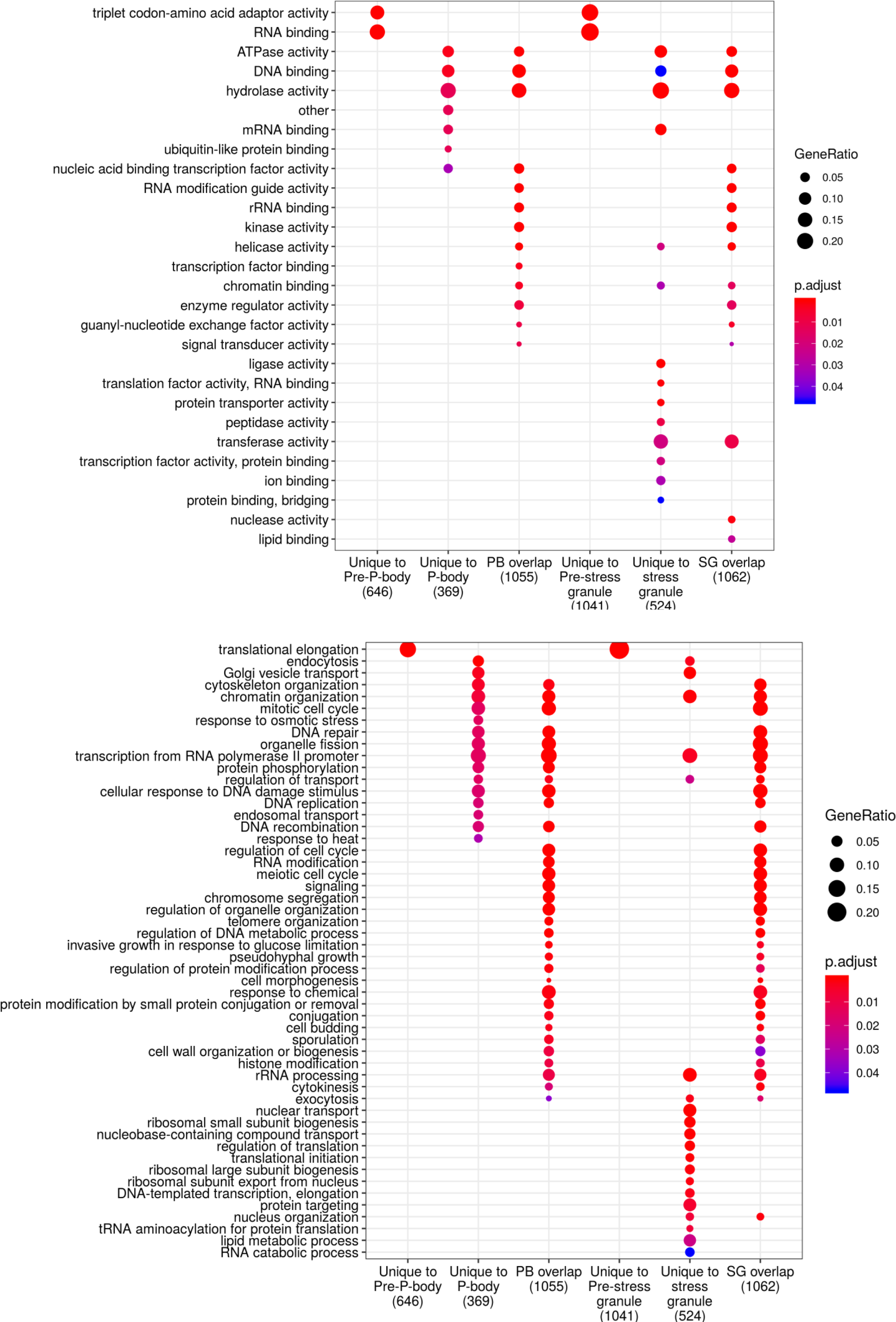
Functional categorisation of mRNAs present in PBs and SGs following glucose depletion.

**Supplementary Fig. 8.**
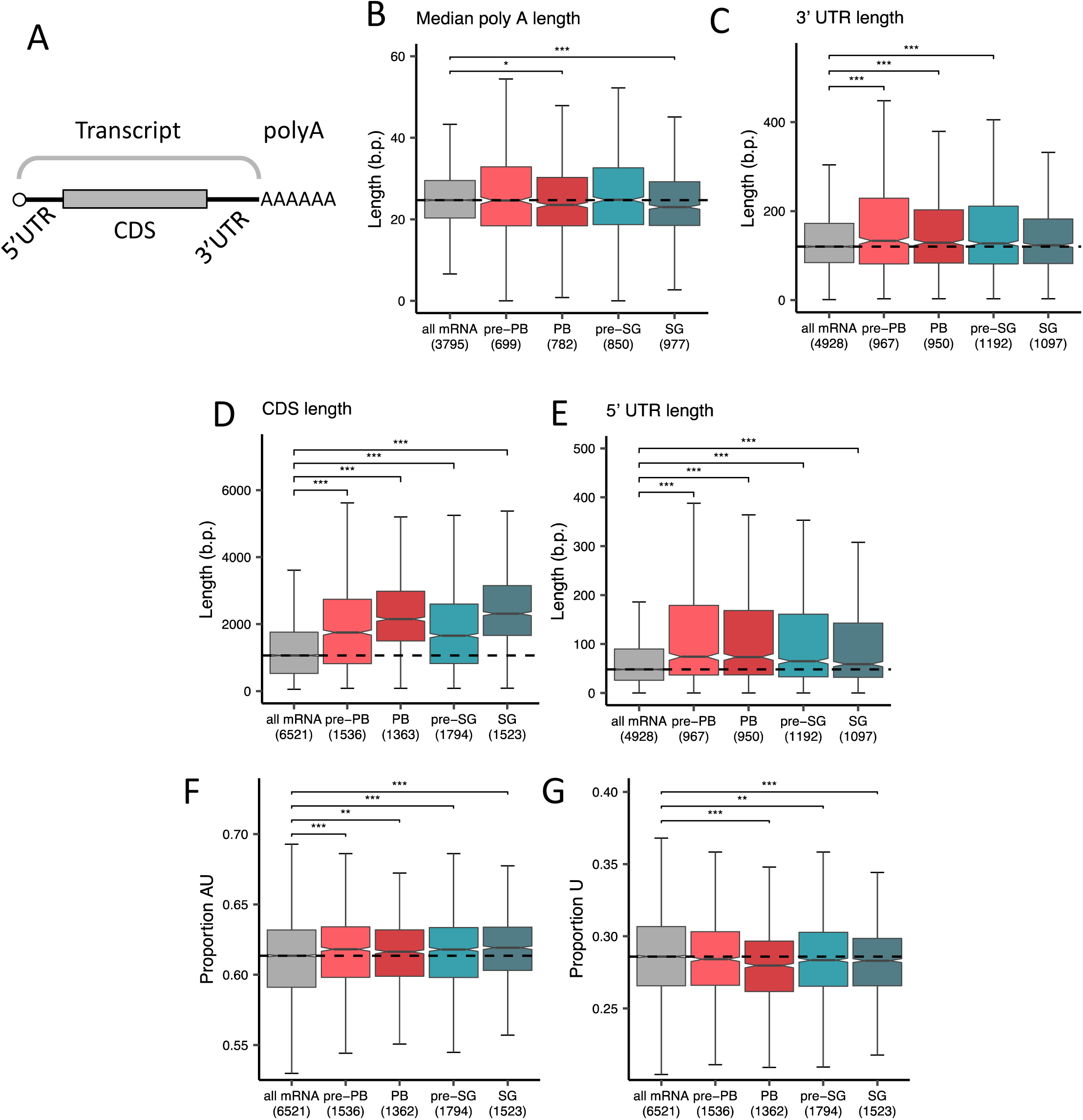
Properties of mRNAs enriched in PBs and SGs. **A.** Schematic of an mRNA illustrating its various components that were individually compared. **B-E.** Box plots comparing the median polyA tail length (**B**), the 5’ UTR (**C**), the CDS (**D**) and the 3’ UTR (**E**) of mRNAs enriched in pre-PBs, PBs, pre-SGs and SGs with all mRNAs from the transcriptome as a background control. **F.** Box plots comparing the AU content of mRNAs enriched in pre-PBs, PBs, pre-SGs and SGs with all mRNAs. **G.** Box plots comparing guanosine content of mRNAs enriched in pre-PBs, PBs, pre-SGs and SGs with all mRNAs.

**Supplementary Fig. 9.**
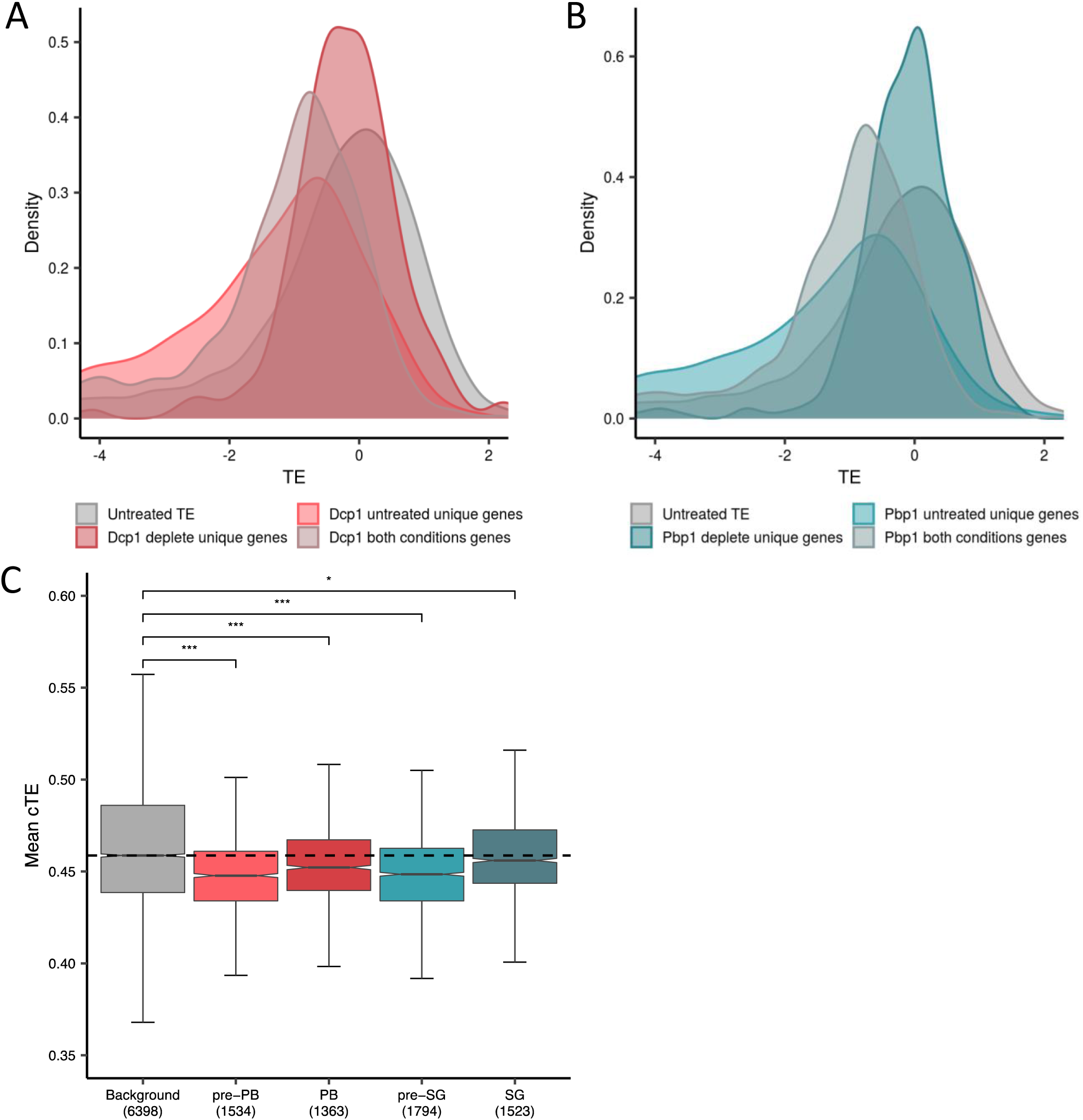
mRNAs enriched in PBs and SGs are translationally repressed. **A.** Comparison of TEs from actively growing cells for mRNAs that are uniquely associated with pre-PBs, PBs or are common to both **B.** Comparison of TEs from actively growing cells for mRNAs that are uniquely associated with pre-SGs, SGs or are common to both **C.** Box plots comparing mean cTEs for mRNAs present in pre-PBs, PBs, pre-SGs and SGs.

**Supplementary Fig. 10.**
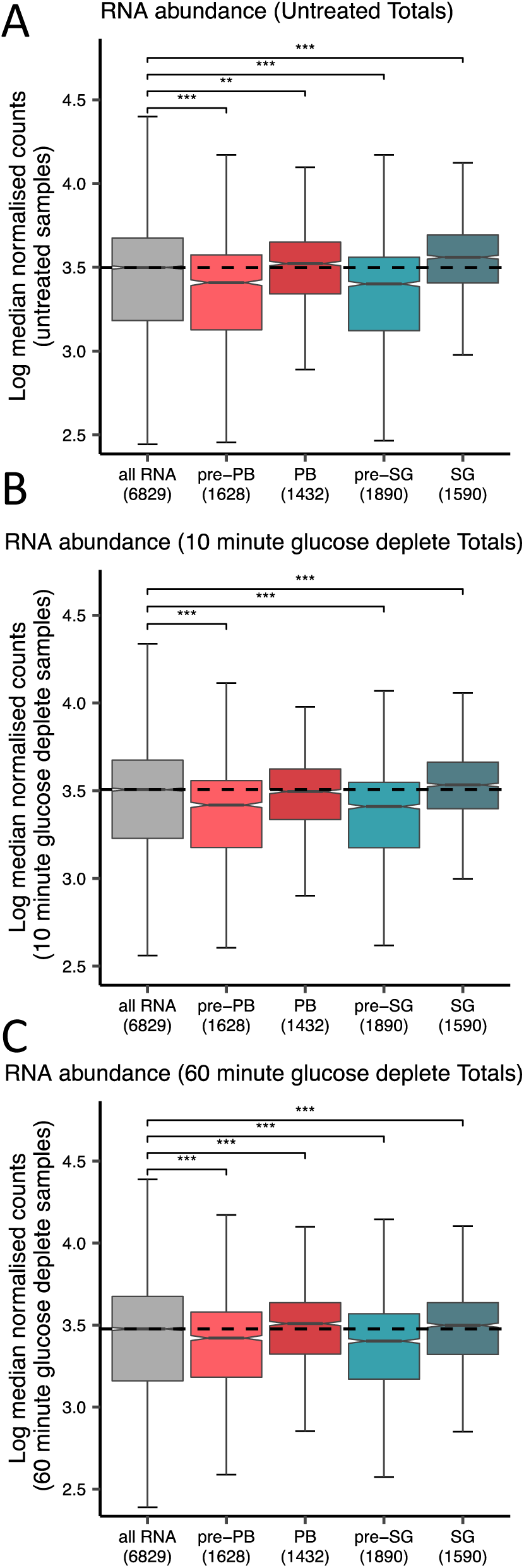
The abundances of mRNAs enriched in PBs and SGs alters between unstressed and conditions of glucose depletion. **A-C.** Box plots comparing mRNA abundances determined using unstressed cells (**A**), cells following 10 minutes glucose depletion (**B**) and cells following 60 minutes glucose depletion (**C**) for those RNAs enriched in pre-PBs, PBs, Pre-SGs and SGs. Background abundance is shown for the transcriptome as a comparison.

